# Selective sweeps in a nutshell; the genomic footprint of rapid insecticide resistance evolution in an insect

**DOI:** 10.1101/828418

**Authors:** Bernarda Calla, Mark Demkovich, Joel P. Siegel, João Paulo Gomes Viana, Kim K.O. Walden, Hugh M. Robertson, May R. Berenbaum

## Abstract

Relatively few genome-wide population studies of field-acquired insecticide resistance have been carried out on agricultural pests. Recently acquired bifenthrin resistance in a population of the navel orangeworm (*Amyelois transitella)*, the main insect pest of almond orchards in California, provided an opportunity to examine the short- and long-term effects of heavy insecticide usage in the population genomic landscape of this species. We re-sequenced the genomes of three contemporary *A. transitella* natural populations differing in bifenthrin resistance status and characterized their population genetics parameters, in the process we detected an exceptionally large selective sweep in all populations. This sweep has virtually no polymorphisms and extends up to 1.3 Mb (spanning 43 genes) in the resistant population. We analyzed the possible causes of this unusually strong population genetic signature, and found genes in the sweep that are associated with DDT and pyrethroid resistance including a cluster of cytochrome P450 coding genes and the gene coding for the small conductance sodium channel “*para*”. Moreover, we found that the sequence along the sweep is nearly identical in the genome assembled from a strain founded in 1966, suggesting that the underpinning for insecticide resistance may have been laid a half-century ago when the California Central Valley experienced massive area-wide applications of DDT for pest control. Our findings are consistent with a scenario whereby insecticide resistance in this species evolved as a stacking of selective pressures that started decades ago and that effectively reduced variation in a region of the genome containing several genes associated with resistance to insecticides with a shared target site and mechanism of action.

## Introduction

Since the rapid post-World War II expansion of the use of synthetic organic insecticides, more than one thousand individual cases of resistance involving more than 500 arthropod species have been documented (Whalon et al., 2008; Sparks and Nauen, 2015). Many of the underlying genetic mechanisms of metabolic resistance and target site insensitivity have been associated with individual genes (Taylor, 1986; Oakeshott et al., 2003). Sufficiently intense pesticide selection, however, can result in acquisition of multiple resistance mechanisms within target pest populations and are associated with multiple genes and larger genomic regions. Studies of selection for insecticide resistance at the genomic scale have been focused mainly on drosophilid model species (Steele et al., 2015; Schmidt et al., 2017; Duneau et al., 2018) or medically important mosquito species (Fouet et al., 2017; Kamdem et al., 2017; Kotsakiozi et al., 2017). Relatively few genome-wide population studies of field-acquired insecticide resistance have been carried out on agricultural pests, a surprising omission given the frequency and economic consequences of resistance evolution in insects.

The primary insect pest of California almonds (*Prunus dulcis)*, a >$ 7 billion industry, is the navel orangeworm (*Amyelois transitella*, Lepidoptera: Pyralidae). In addition to almonds, the highly polyphagous *A. transitella* also causes significant losses in other nut and fruit tree crops, including pistachios (*Pistacia vera)*, walnuts (*Juglans* spp.), and figs (*Ficus carica*). Nut crops are vulnerable to oviposition by *A. transitella* adults and damage by larvae when kernels are exposed at hull split. Beyond this direct damage, the navel orangeworm is a facultative mutualist of the fungal pathogen *Aspergillus flavus* and, by serving as a vector, can increase the likelihood of contamination by aflatoxins, reducing crop quality and marketability (Palumbo et al., 2014). Adding to its potential for causing economic losses is the fact that, in California, *A. transitella* is multivoltine, with up to four generations per year depending on weather.

Insecticides have been used intensively in almond orchards to protect them from the navel orangeworm, and these are generally applied at hull split to reduce feeding damage, and up to three times over the course of the growing season. Although insecticides approved for use include representatives from five structural classes, pyrethroids have been widely used due to their low cost and high efficacy. Prior to 2013, insecticide resistance was not known to occur in navel orangeworm populations; in 2013, however, ten-fold resistance to bifenthrin was documented in populations in Kern County (Demkovich et al., 2015).

Taking advantage of the recent sequencing of the *A. transitella* genome (http://i5k.github.io/), we used a population genomics approach to locate regions of the genome under insecticide selection; We evaluated nucleotide diversity and genomic differentiation across windows of the genome in three modern populations of this species. Additionally, we characterized the population genomic parameters in families of genes whose members are often involved in xenobiotic detoxification in Lepidoptera. In the process, we detected an extremely large selective sweep across all of the sequenced populations. In analyzing the mutations in genes along the sweep and comparing them with variants in the reference genome, we discovered that the reference genome, from a laboratory strain founded in 1966, has an almost identical sequence along ∼0.6 Mb of the sweep. We then review patterns of pesticide use in the Central Valley to understand the origins of this selection signature.

## Materials and Methods

### Genome re-sequencing and identification of genomic regions with positive selection hallmarks (Pool-seq)

The ALM and FIG strains of *A. transitella* were collected from orchards in Madera County as larvae (JPS, USDA-ARS, Parlier, CA) in 2016. Larvae from the pyrethroid-resistant strain R347 were collected from almond orchards in Kern County and sent to us by Brad Higbee (Trécé) in 2016. All three populations were maintained in an incubator at University of Illinois at Urbana-Champaign at temperatures of 28 ± 4°C and photoperiod of 16:8 (L:D) h cycle, reared until adulthood, and separated by sex before freezing at −80°C. Genomic DNA was extracted from the heads of 100 adult moths (equal sex ratios) Insects were ground in liquid nitrogen, lysed overnight with SDS and Proteinase K, treated with RNase A, and centrifuged in a high-salt solution to precipitate proteins. The DNA was precipitated with ethanol, re-suspended in 10 mM Tris pH 8, and evaluated quantitatively and qualitatively with a Qubit fluorometer (Thermo Fisher Scientific, USA) and checked for degradation on an agarose gel. Subsequently, 2.5 µg of male head DNA were combined with 2.5 µg of female head DNA into a single tube for each of the three strains. Shotgun genomic libraries were prepared with the Hyper Library construction kit (Kapa Biosystems, Wilmington, MA) from equimolar-pooled DNA samples from each of the three populations. Libraries were quantitated with qPCR and paired-end sequenced for 150 bases-long reads on one lane of the Illumina HiSeq 4000 (Illumina, San Diego, CA). Library construction and sequencing were carried out at the W.M. Keck Center of the Roy J. Carver Biotechnology Center at the University of Illinois at Urbana-Champaign.

### Alignment, SNP calling and calculation of population genomic signtures

Reads were trimmed for residual adapters and quality using trimmomatic v. 0.32 (Bolger et al., 2014). The trimmed reads were aligned to the *A. transitella* reference genome (NCBI accession ASM118610v1) using bwa mem v. 0.7.12-r1039 with paired-end mode (Li, 2013). The resulting SAM files were sorted and optical- and PCR-derived sequencing duplicates were marked and removed using Picard v.1.48 (Broad institute, http://broadinstitute.github.io/picard/). Low-quality alignments (mapping quality score < 20), improper pairs and un-mapped mate reads were removed using SAMtools v.1.7 (Li et al., 2009). A pileup file was created for each separate library using the mpileup command from SAMtools. Indels were identified and separated from the main pileup files. The three pileup libraries were then sub-sampled for uniform SNP coverage of reads using ‘identify-genomic-indel-regions.pl’ and ‘subsample-pileup.pl’ scripts from the Popoolation v.1.2.2 package utilizing the max-coverage =100 parameter (Kofler et al., 2011a). The subsampling was done to prevent biases in population genomic metrics affected by sequencing errors and copy number variation that could skew coverage in affected regions. Tajima’s π was calculated for each of the strains (libraries) using 5 kb-long windows and 5 kb-long steps using the ‘Variance-sliding.pl’ script from Popoolation. On a separate pipeline, a multiple-pileup file was created that incorporated the three trimmed and quality-filtered mapped libraries, indels were removed as described and libraries were synchronized using ‘mpileup2sync.jar’ from Pooplation2 software (v.1.201) (Kofler et al., 2011b). The 3-populations pileup file was subsampled for uniform coverage and pair-wise *F*_*ST*_ values were calculated using the ‘fst-sliding.pl’ script from Popoolation2 on a per-SNP basis.

For pooled DNA sequencing, *F*_*ST*_ values might present bias if the sample is too small or if non-equimolar amounts of DNA for each of the individuals from each population were used. Bias can also be introduced during the PCR amplification step before sequencing (Cutler and Jensen, 2010). Even though *F*_*ST*_ calculation in Popoolation2 implements a bias correction (Kofler et al., 2011b), the estimates are still deemed biased according to Hivert et al. (Hivert et al., 2018). For that reason, to validate our *F*_*ST*_ estimates, the synchronized file with data from the three populations obtained using ‘mpileup2sync.jar’ from Pooplation2 software (see above section), was converted to a pooldata object for the “Poolfstat” v. 1.0.0 R package (https://cran.r-project.org/web/packages/poolfstat). Both methods had similar results, but Popoolation2 does not report overall *F*_*ST*_ between populations; accordingly, we report results from both packages. ‘Pcadapt’ v. 1.1 (50) was used to detect the outlier SNPs based on Principal Component Analysis (PCA). In PCAdapt, z-scores were calculated based on the original set of SNPs using K = 2 principal components. Outliers were then identified on the z-scores vector using Mahalanobis distances. The distances were transformed into *P*-values assuming a chi-square distribution with K degrees of freedom (Luu et al., 2017).

### Sanger sequencing the para locus in laboratory strains

For SPIRL-1996, we obtained genetic material from frozen whole-body fifth instar larvae, as the strain is no longer available as a laboratory colony. DNA was extracted using an E.Z.N.A.® insect DNA kit (Omega Bio-tek, Norcross, GA) according to the manufacturer’s instructions. For the CPQ strain, we utilized existing midgut cDNA sampled from ten different individuals; The available CPQ cDNA was previously prepared from total RNA derived from midguts of fifth-instar larvae fed on semi-synthetic diet using a Protoscript II kit (NEB, Ipswich, MA). PCR was carried out on both strains using primers designed to flank the region of the *kdr* mutation in the para gene (Forward 5’-ACCAAGGTGGAACTTCACAGAT −3’ Reverse 5’-AGCAATTTCAAGAAGTCAGCAACA −3’). PCR amplicons were sequenced and sequences were aligned to the reference to verify the presence or absence of the mutation.

### Insecticide bioassays

To establish the median-lethal concentrations (LC_50_) for bifenthrin and DDT in the sequenced strains, we used feeding assays with semi-synthetic artificial diet (Waldbauer et al., 1984). Bifenthrin (Chem Service Inc., West Chester, PA), or DDT (Sigma-Aldrich Co., St. Louis, MO) was stirred into the diet at different concentrations for each strain and poured into separate 1-oz (28 ml) cups to set. Treatments and concentrations were: bifenthrin in methanol – ALM: 2 ppm, 5 ppm, 10 ppm, 12 ppm, 15 ppm, 24 ppm; DDT – ALM: 50 ppm, 100 ppm, 200 ppm, 300 ppm, 400 ppm; bifenthrin – R347: 8 ppm, 16 ppm, 24 ppm, 48 ppm, 75 ppm; DDT – R347: 50 ppm, 100 ppm, 200 ppm, 300 ppm, 400 ppm; DDT – CPQ: 10 ppm, 20 ppm, 35 ppm, 50 ppm, 75 ppm, 100 ppm. Four neonates were transferred with a soft brush into each plastic cup containing bifenthrin or methanol as the solvent control. Twenty larvae from each strain were exposed to their respective bifenthrin or DDT concentrations and each assay was replicated three times per concentration. Neonate mortality on diets was assessed after 48 h and scored according to a movement response after being touched by a soft brush. Probit analysis (SPSS version 22, SPSS Inc., Chicago, IL) was applied to identify median-lethal concentrations (LC_50_). Differences between populations were considered significant if their respective 95% confidence intervals in the Probit analysis did not overlap. We were unable to perform these assays with the FIG strain because we did not establish a colony from the population used for sequencing; however, for the purposes of comparison we cited the bifenthrin LC_50_ established in our laboratory by Bagchi et al. (Bagchi et al., 2016).

### Sequencing the para locus in laboratory strains

DNA was extracted using ten frozen whole-body fifth instar larvae from the reference genome strain SPIRL-1966, and ten midguts of fifth instar larvae of the CPQ strain that fed on semi-synthetic artificial diet (Waldbauer et al., 1984). Extractions were done using an E.Z.N.A.® insect DNA kit (Omega Bio-tek, Norcross, GA) according to manufacturer’s instructions. Primers were designed using Primer3 (Untergasser et al., 2012) as implemented in Geneious software (Kearse et al., 2012) to sequence the region of the *para* gene containing the *kdr* mutation (Forward 5’-ACCAAGGTGGAACTTCACAGAT −3’ Reverse 5’-AGCAATTTCAAGAAGTCAGCAACA −3’). Amplicons were Sanger sequenced and trace analysis and alignments were performed in Geneious.

### Obtaining pesticide application data

Records of pyrethroid use were accessed through the California Department of Pesticide Regulation (CDPR) - pesticide use annual reports from 1990-2016. Total bifenthrin use in almond orchards was analyzed in Kern County, Madera County, and statewide based on number of applications, pounds of active ingredient, and acres treated from 2006-2016. We also examined records of all pyrethroids applied in almonds from 2000-2016 and compared bifenthrin use relative to all registered pyrethroids by pounds of active ingredient and acres treated.

## Results

### Genome re-sequencing and population genomics parameters

One hundred individuals from each of three navel orangeworm populations from the Central Valley were sampled and sequenced in pools. In addition to the resistant population in Kern County (R347), two populations were collected in Madera County: one from almond orchards (ALM), and one from figs (FIG). Both populations displayed field susceptibility to bifenthrin at the time of collection. The sequencing produced more than 700 million 150 nt-long paired reads with an average Phred quality score higher than 30 throughout each read and resulted in roughly 82X raw coverage of the 409-Mb *A. transitella* genome for each of the strains sequenced (Table S1).

The reads were mapped to our *A. transitella* reference genome, which was sequenced from a laboratory strain (SPIRL-1966). After mapping, filtering, and subsampling reads, the total number of called SNPs for the ALM population was 10,592,326, for FIG was 10,303,426 and for R347 was 9,704,346. The nucleotide diversity, measured as Tajima’s π across non-overlapping 5kb-long windows for each of the populations, was on average 0.0179, 0.0175, and 0.0171, for ALM, FIG, and R347, respectively. An unusually large region displaying substantially reduced genetic diversity across the three populations was readily detected by screening Tajima’s π values that were calculated with 5kb non-overlapping windows across the genome (Table 1). We narrowed down this region of low nucleotide diversity to the region starting around 2.92 Mb in Scaffold NW_013535362.1 and extending up to the base at approximately 3.8 Mb in the ALM and FIG populations and up to the base at approximately 4.32 Mb in the R347 line. The region of low nucleotide diversity that spans the three populations encompasses 38 genes. In the resistant line, the total selective sweep extends to up to 43 genes (Figure 1, Panel A). An examination of the read alignments to the reference strain SPIRL-1966 shows that the nucleotide sequence of the reference is nearly identical to that of the three other populations along the hard sweep region. In addition, the reference genome sequence is also nearly identical to that of R347 in the regions flanking the sweep (Figure S1).

**Table 1.** Single nucleotide polymorphisms and Tajima’s π calculated on 5Kb non-overlapping windows across the whole genome in three populations of *Amyelois transitella*, ALM and FIG susceptible, R347 resistant.

|  | ALM | FIG | R347 |
| --- | --- | --- | --- |
| **number of SNPs** | 10592326 | 10303426 | 9704346 |
| **Tajima's $\pi$**<br>**(genome-wide average)** | 0.17911 | 0.017517 | 0.017089 |
| **Minimum value of $\pi$**<br>**(scaffold)** | 0.000018987<br>(NW_013535362) | 0.000047803<br>(NW_013535362) | 0.000000000<br>(NW_013535362) |
| **Maximum value of $\pi$**<br>**(scaffold)** | 0.039661659<br>(NW_013535371.1) | 0.040058152<br>(NW_013535371.1) | 0.039509560<br>(NW_013535951.1) |
| **In NW_013535362.1 only (calculated on coding regions)** |  |  |  |
| **Tajima's $\pi$ average** | 0.0148181 | 0.0142969 | 0.0131416 |

**Figure 1.**
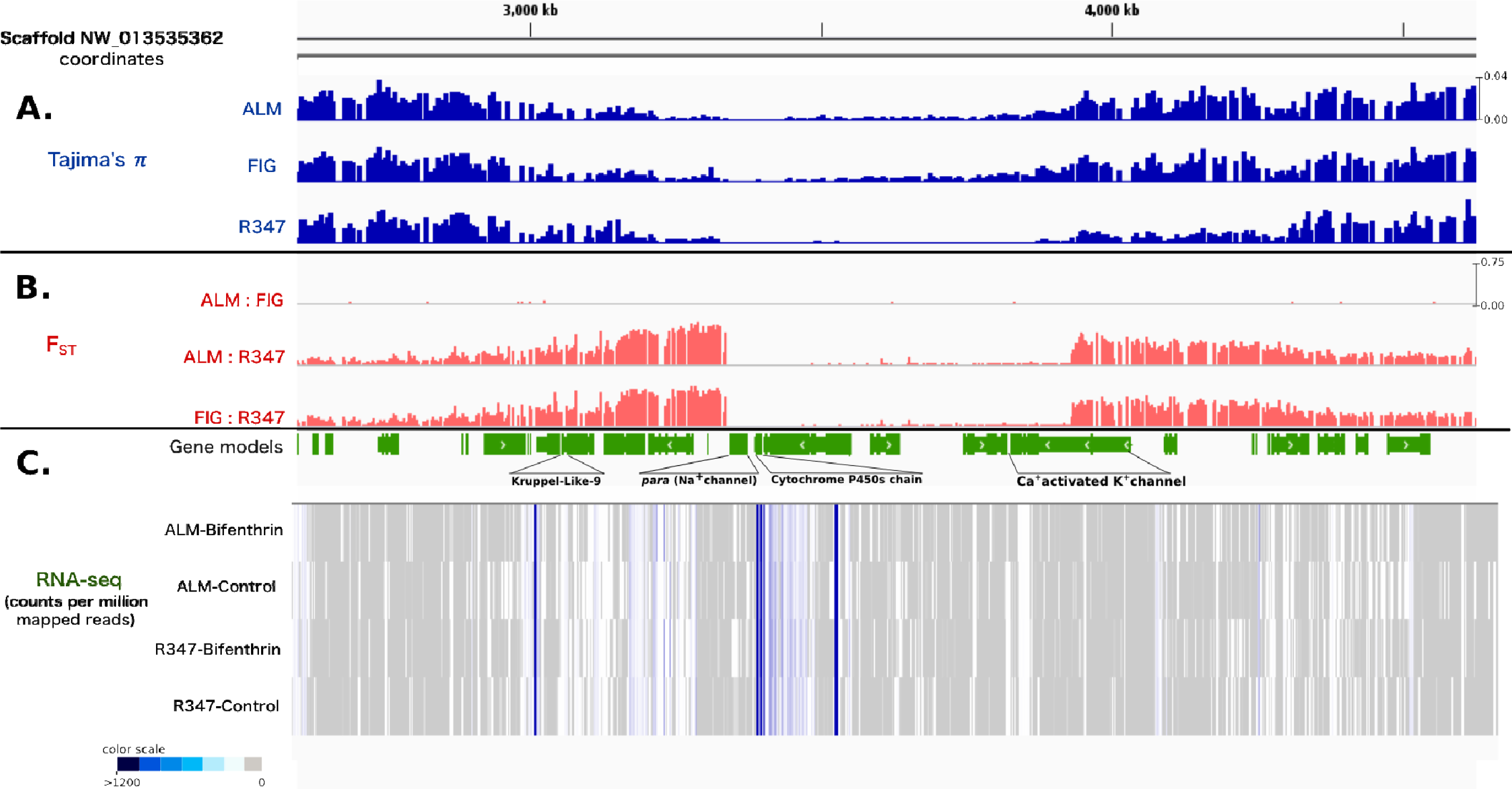
Selective sweep region in Scaffold NW_013535362.1 of the *Amyelois transitella* genome. (A) Nucleotide diversity measured as Tajimas π in two bifenthrin susceptible populations (ALM and FIG), and in one bifenthrin resistance population (R347). (B) Pairwise FST estimates for the three populations and (C) RNA-seq data for ALM and R347 fifth-instar individuals exposed to bifenthrin in counts per million mapped reads along the sweep region.

Selective sweeps are the result of beneficial mutations that rapidly became fixed in the population and carried along neighboring variants, resulting in a local loss of heterozygosity; in a complete hard sweep, heterozygosity reaches zero at or very close to the causative mutation (Burke et al., 2010; Burke, 2012). To further narrow down the causative mutation that could have generated the observed pattern, the Tajima’s π metric was re-calculated on a gene-by-gene basis (where the length of the window to account for variable nucleotides is equal to the length of the gene) on the predicted gene models across the entire NW_013535362 scaffold. Of the 190 annotated gene models, 170 had enough sequencing coverage to calculate nucleotide diversity at the set cut-offs. In the ALM population, nucleotide diversity reached zero (π = 0) at the *protein kish-A* gene (XM_013328236.1), with no SNPs detected along this coding sequence, followed by the voltage-gated sodium channel “*para*” (XM_013328250.1), with a single SNP. In the FIG population, the lowest π value (0.0001) was also found in *para*, followed by the gene encoding the cytochrome P450 CYP6B56 (XM_013328369.1). In the R347 strain, π was zero at the gene coding for CYP6B56, followed by the gene coding for “protein Ariadne”, a putative RING-type transcription factor (Table 2). The coverage and number of SNPs for each of the genes in the scaffold is detailed in Table S2).

**Table 2.**
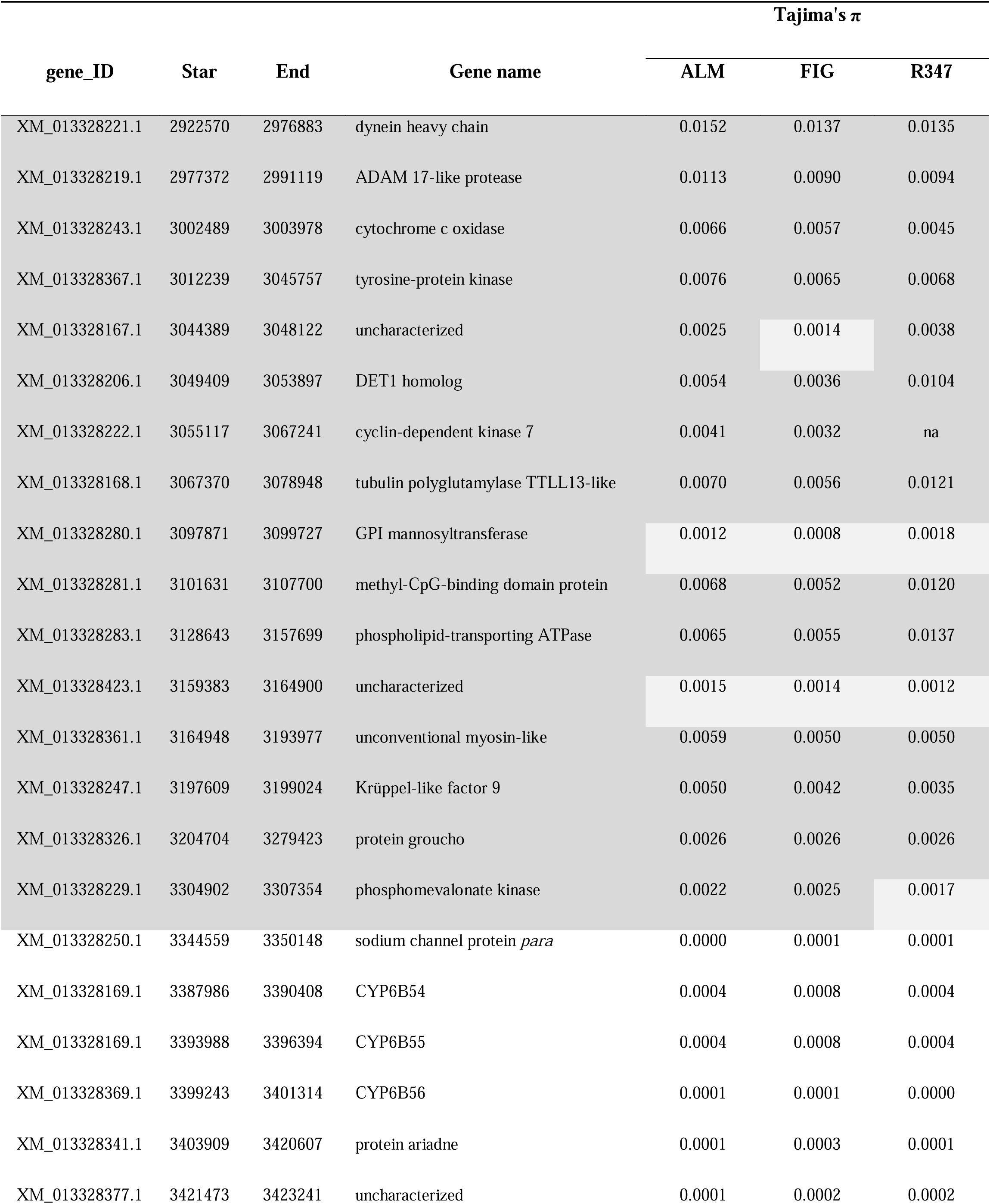
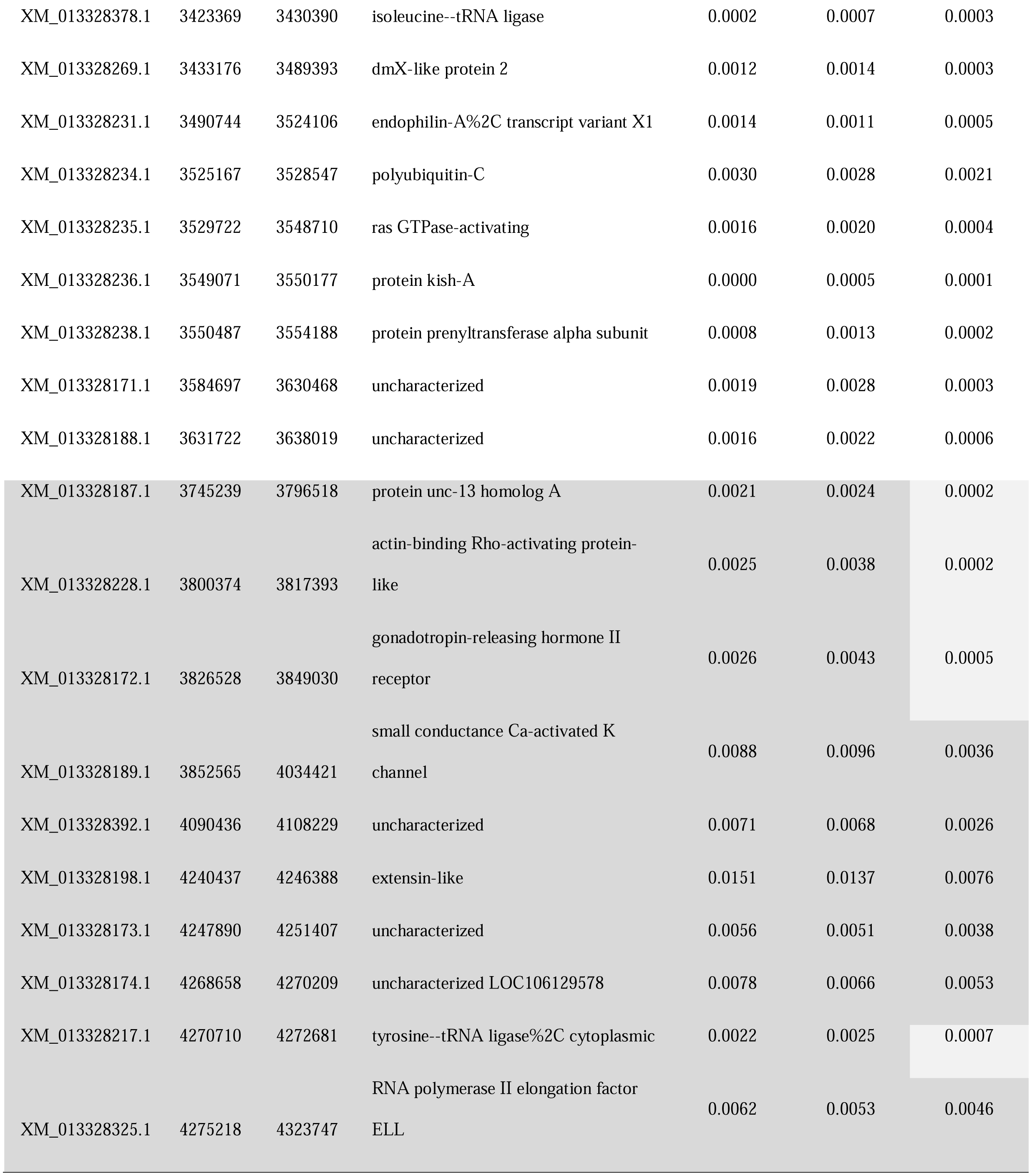
Nucleotide diversity estimates for genes in the sweep region (Scaffold NW_01353536). Only genes with enough read coverage to measure Tajima’s π are shown (41 out of 44). (Dark gray cells = π >0.02, light gray and white cells π <0.002, white cells = defined hard sweep region in all three populations).

We then manually screened the few nucleotide variations along the region where all three populations have near zero π values. Only two genes had non-silent mutations in all or in a portion of the reads covering the position across the three populations. One of these was the *para* gene (which also showed Tajima’s π values equal to zero), with the mutation L934F. All three populations carried the mutation in 100% of the sequenced reads covering the position, in contrast with the reference genome (Figure 2, panel A). This mutation corresponds to the known *kdr* (‘knock down resistance’) mutation that confers target site resistance to DDT and pyrethroids (Williamson et al., 1996) in multiple insect species (Miyazaki et al., 1996; Pittendrigh et al., 1997; Bass et al., 2007; Haddi et al., 2012; Dong et al., 2014). Although the reference genome shows nearly identical nucleotide sequence along *para*, it does not have the *kdr* mutation. In the absence of population level data, we confirmed that the reference strain did not carry the *kdr* mutation by re-sequencing the region flanking this mutation on ten SPIRL-1966 individuals collected in 2012 and preserved in our laboratory (Figure S2). A second gene encoding a *Krüppel-like-9* factor (KL-9) had non-silent mutations segregating in 82% and 91% of the sequenced reads from ALM and FIG, respectively, and in only 9% of the reads in the resistant R347. KL-9 overlaps the region where all three populations have low diversity and the region where only the R347 strain has reduced π values (Figure 1A, Figure 2B, and Table S3).

**Figure 2.**
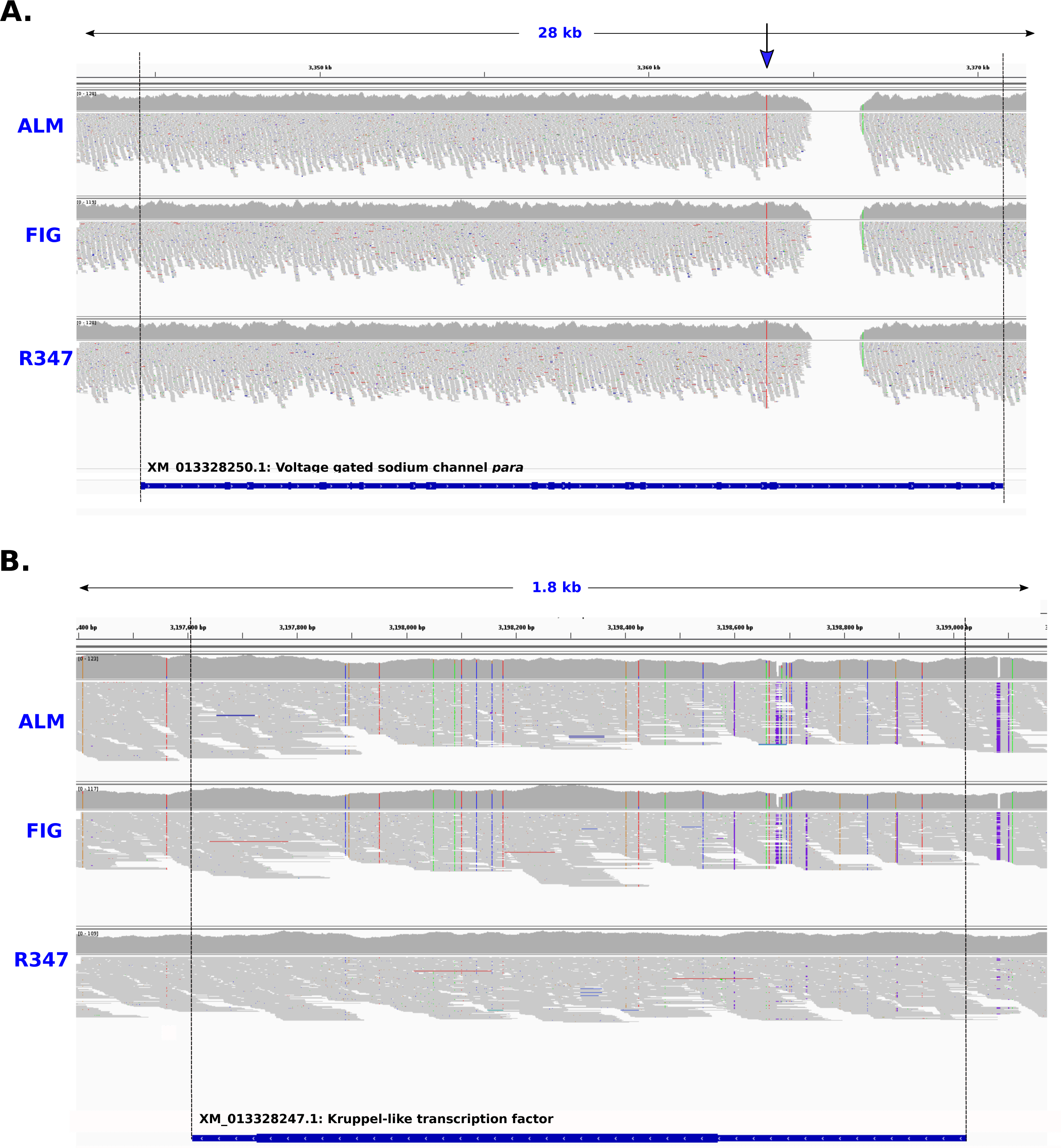
Alignment of reads for the three sequenced populations to the SPIRL-1966 reference genome in the voltage-gated sodium channel *para* (A), and in the Krüppel-like transcription factor (B), both located in the sweep region in Scaffold NW_013535362.1 of the *Amyelois transitella* genome. Gray color signifies exact base match, colored lines are mutations relative to the reference genome. The arrow in (A) shows the only mutation found in para which correspond to the *kdr* mutation that confers resistance to DDT and pyrethroids.

We then calculated nucleotide diversity in the mRNA sequences of members of four detoxification gene families whose members might be involved in insecticide resistance -- cytochrome P450s, glutathione-S-transferases (GSTs), ABC transporters, and carboxylesterases (COEs) -- across the whole genome. The calculation of Tajima’s π in this case used the gene length as window size. In addition to the three P450 genes located in the selective sweep, 57 other P450 genes had enough read coverage to calculate nucleotide diversity according to our set cut-offs. The R347 population showed a low π (i.e., less than half the median of the three populations in the gene) in CYP6AW1 relative to ALM and FIG and relative to the rest of P450s in the set, suggesting possible local positive selection on this gene in the resistant line (Figure S3 A). Among the ABC transporters, 29 of 56 had enough read coverage in at least two of the populations to calculate π but none of them had nucleotide diversity π below half of the median of the three populations, with the lowest π values in the gene coding for Atra_ABC-C6 (XM_013339562.1). Among the GSTs, 18 had enough SNP coverage to calculate a π, but none was below the median of the three population and *Atra*_GSTω1 (XM_013335304.1) had the lowest π across all GSTs in the three populations (Figure S3 B). Forty-two of 66 COEs had enough SNP coverage, where a clade B α-esterase (XM_013334043.1) has the lowest nucleotide diversity in the three populations, and no COE had significantly lower π when compared across populations (Figure S3 C).

Genetic differentiation measured as *F*_ST_ between populations showed that ALM and FIG are probably indistinguishable as separate strains (*F*_ST_ −0.004, as negative *F*_ST_ values are effectively zero). The *F*_ST_ between ALM and R347 was 0.0289, and between FIG and R347 was 0.0290. These values are highly reflective of the demographic origins of each population (i.e., Madera County, where ALM and FIG were sampled vs. Kern County, the source for the R347 strain). Genes controlling traits that differ between populations might present large differences in allele frequencies, thereby generating higher *F*_ST_ values comparable with average *F*_ST_ across the genome (Myles et al., 2008). We scanned for outlier loci showing SNPs with large differences in allele frequency by calculating *F*_ST_ on a by-SNP basis utilizing two different methods (see Materials and Methods). The SNPs with top 1% *F*_ST_ values were concentrated in the regions flanking the selective sweep (Figure 1B and Figure S4). These high *F*_ST_ values were all found in pairwise comparisons with the resistant genotype (i.e. ALM vs. R347 or FIG vs. R347).

The *F*_ST_ method may not account for unexpected population structure and requires that each sequenced individual is assigned to a single population, an assumption that might not be valid. To verify our findings by overcoming the limitations of *F*_ST_, we utilized an approach based on principal component analysis (PCA) on the calculated major allele frequencies (Luu et al., 2017), under the assumption that SNPs that are highly correlated with underlying population structure are candidates for local adaptation. We screened the top 20 differentiated SNPs (FDR-corrected p-value <0.07, k=2). Of those, only 2 SNPs were differentiated between the resistant and susceptible populations: one in scaffold NW_013535509.1, in an intergenic region near the ecdysteroid kinase gene cluster and a second in the NW_013535362.1 scaffold, just downstream of the selective sweep in position ∼4.1Mb (Table S4, Fig 3). To assess whether the high *F*_ST_ outliers found in scaffold NW_013535362.1 are responsible for the overall genetic differentiation between populations, we re-calculated the overall pairwise *F*_ST_ values after removing the region between position 2.95 Mb and 3.93 Mb in NW_013535362.1. The resulting *F*_ST_ values were almost identical to the ones obtained with the full genomes, indicating that, other than the region surrounding the selective sweep, these populations are likely differentiated as a consequence of local (i.e., geographically restricted) selection, including population bottlenecks, genetic drift, or migration.

**Figure 3.**
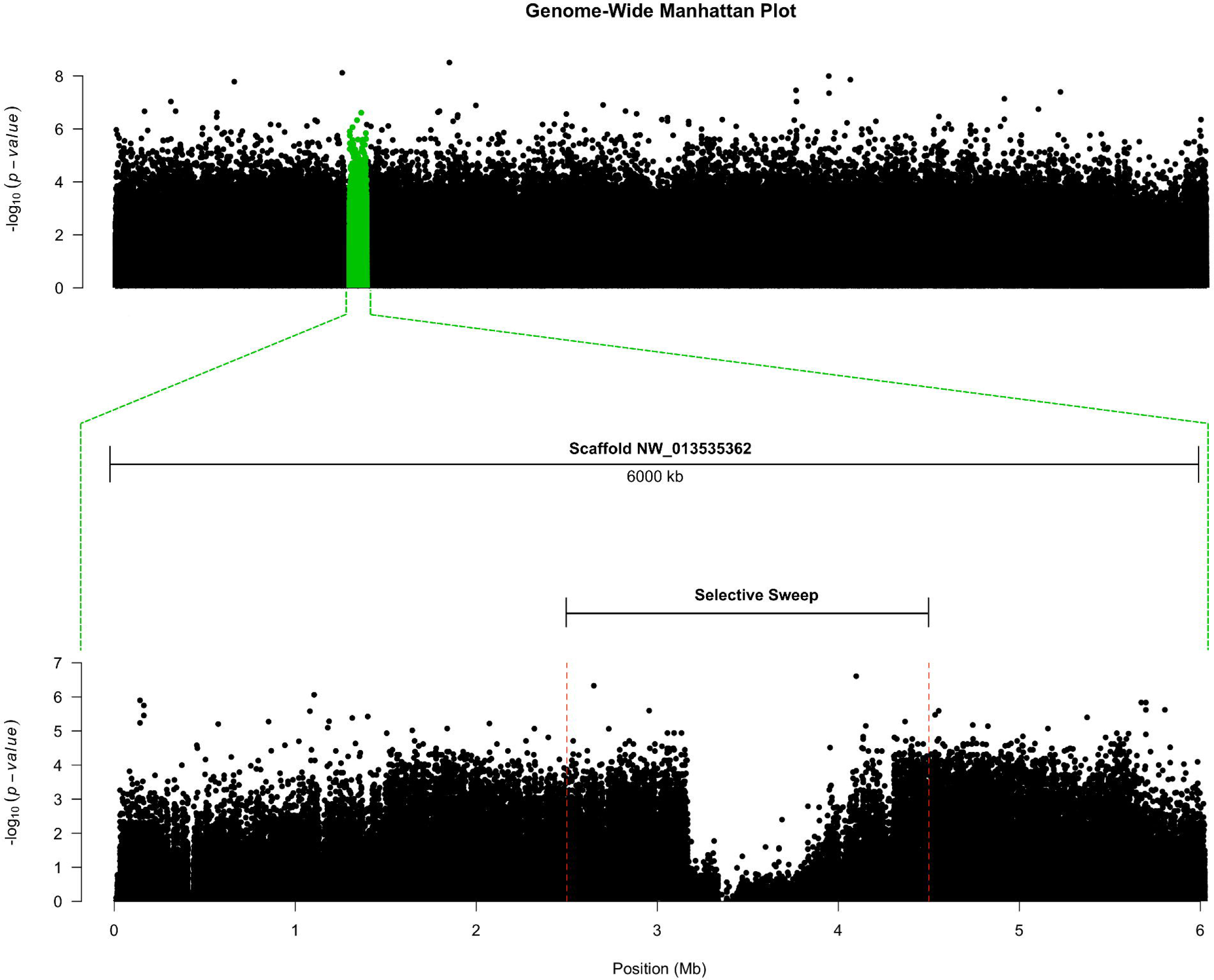
Manhattan plot for the tree tested populations of *Amyelois transitella*. The plot is based on the *p-values* of Mahalanobis distances between calculated z-scores derived from regressing each SNP by 2 principal components (*k=2*). For details see Materials and Methods.

Given the size and the strength of the signature of selection that we identified, we hypothesized that genes other than *para* could be involved in insecticide resistance. Of the genes in the sweep, the chain of three cytochrome P450 genes (*CYP6B54, CYP6B55* and *CYP6B56*) are strong candidates, given that genes in this subfamily have been implicated in pyrethroid resistance in other species (Wang and Hobbs, 1995; Ranasinghe and Hobbs, 1998; Zhang et al., 2010). Tandem clusters of P450s are likely recent duplications that have persisted due to their adaptive value. We compared syntenic regions of *A. transitella* and *Bombyx mori* (the domestic silkworm) and found only one *B. mori* P450 (CYP6B29) adjacent to *para*. In the Old World bollworm *Helicoverpa armigera*, however, there is also a chain of three CYP6B-subfamily genes just adjacent *para*, these same P450s that were previously associated with pyrethroid resistance in this species (Wang and Hobbs, 1995; Ranasinghe and Hobbs, 1998) (Figure 4). Available RNA-seq data (NCBI accession: PRJNA548705), show that transcripts for CYP6AB54, CYP6AEB55, and CYP6AB56 are highly constitutively expressed in samples from both ALM and R347 populations (i.e., up to tenfold higher than the average counts per million in the region); in bifenthrin-treated individuals from the R347 resistant population, CYP6AB56 transcript levels are upregulated up to 1.6 fold relative to non-treated individuals (Table S5 and Figure 1C).

**Figure 4.**
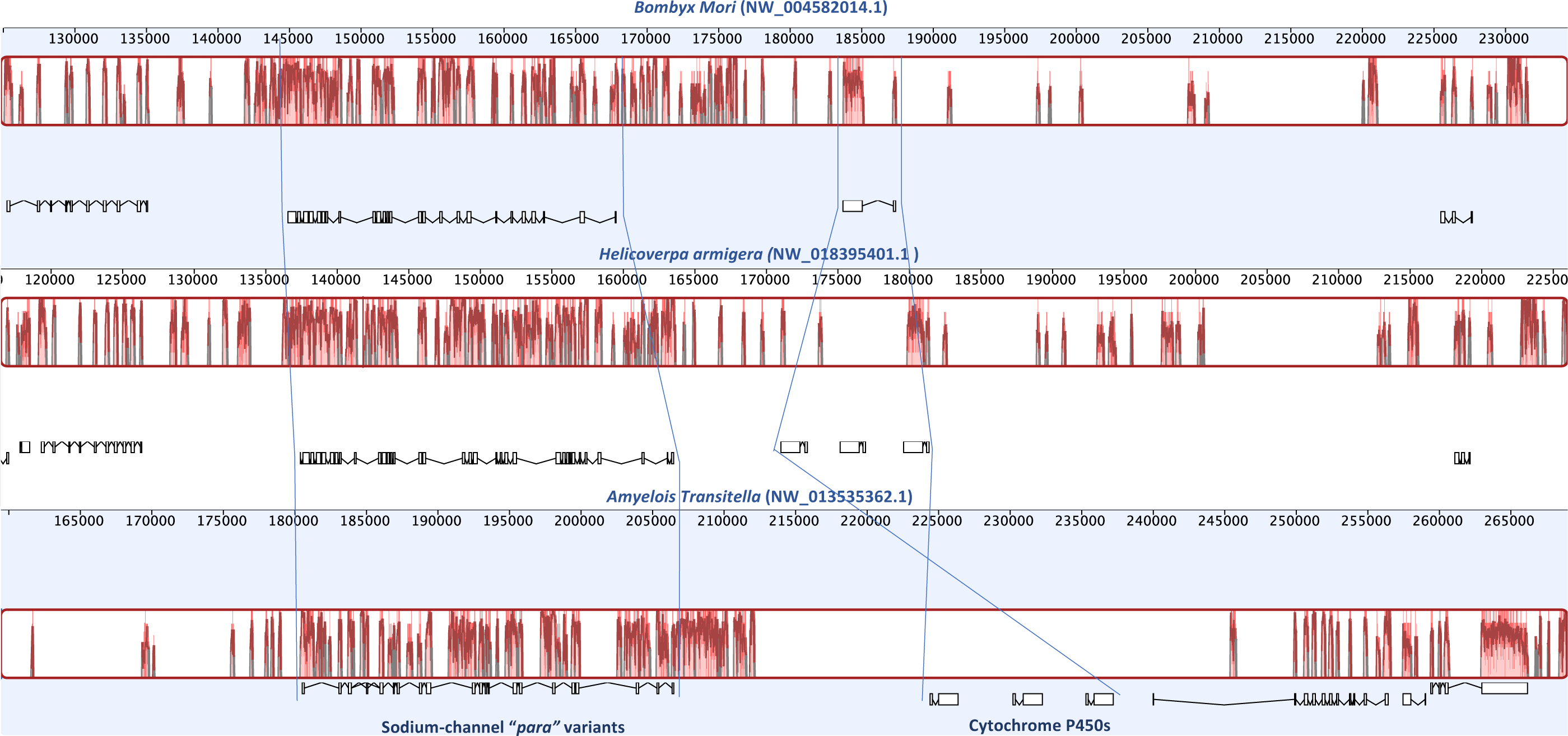
Syntenic regions between *Bombyx mori, Amyelois transitella* and *Helicoverpa armigera*. The region shows the *para* sodium channel gene models next to the cytochrome P450s from the CYP6B subfamily. Both *A. transitella* and *H. armigera* are pest species that have been exposed to heavy insecticide usage.

### Bioassays and analysis of historical data on insecticide usage

To determine the functional significance of the identified genomic signatures, we compared levels of resistance to bifenthrin and DDT in ALM and R347 with CPQ, a strain derived from the original SPIRL-1996 line that lacks the *kdr* mutation. The CPQ strain was the most susceptible to bifenthrin, with an LC_50_ value of 0.38 ppm; in contrast, the R347 strain displayed the highest level of resistance, with a LC_50_ value of 24.27 ppm, confirming field observations of insecticide failure. The LC_50_ of ALM was 7.45, comparable to that of FIG reported by Bagchi et al. (Bagchi et al., 2016). There was no difference in LC_50_ for DDT between the ALM and the R347 populations, both displaying relatively high resistance to DDT (LC_50_=259.85 and 310.33 ppm, respectively) compared to CPQ (LC_50_ =25.32 ppm) (Table 3). These values are suggestive of a direct association between the presence of *kdr* and DDT resistance but not between *kdr* and bifenthrin resistance.

**Table 3.** Comparison of toxicity of bifenthrin and DDT in four strains of navel orangeworm in relation to the presence or absence of the *kdr* mutation.

|  | **Bifenthrin 48 h LC<sub>50</sub>** | **DDT 48 h LC<sub>50</sub>** | ***kdr*** |
| --- | --- | --- | --- |
| **Population** | **(95% CI)** | **(95% CI)** | **mutation** |
| **ALM** | 7.45 (5.90 - 9.64) | 259.85 (216.71 - 326.0) | YES |
| **FIG** | 7.20 (5.35 - 11.08)<sup>1</sup> | ---- | YES |
| **R347** | 24.27 (18.19 - 33.09) | 310.33 (249.37 - 424.29) | YES |
| **CPQ** | 0.38 (0.31 - 0.46)<sup>1</sup> | 25.32 (19.73 - 30.84) | NO |

<sup>1</sup> LC<sub>50</sub> identified in Bagchi et al. (2016)

To gauge the level of selective pressure exerted by bifenthrin usage in the sampled populations, we reviewed historical patterns pesticide usage using data from the California Department of Pesticide Regulation (CDPR) annual reports from 1990 to 2017. Statewide, the use of bifenthrin increased 7.3-fold by pounds applied and 4.5-fold by treated acres from 2007 to 2013 (Figure S5). During this period, Kern County applications in almonds increased 3.5-fold by pounds of bifenthrin applied and 4.4-fold by treated acres, while Madera County applications increased 10.1-fold in pounds applied and 5.2-fold by treated acres. Applications of bifenthrin per pound, however, were higher in Kern County throughout this period, except in 2012 (Figure 5). From 2009 to 2013, the number of registered products containing bifenthrin that were applied in almond orchards increased from 1 to 13 (Table S6). By 2017, 19 products containing bifenthrin were used to treat almond orchards.

**Figure 5.**
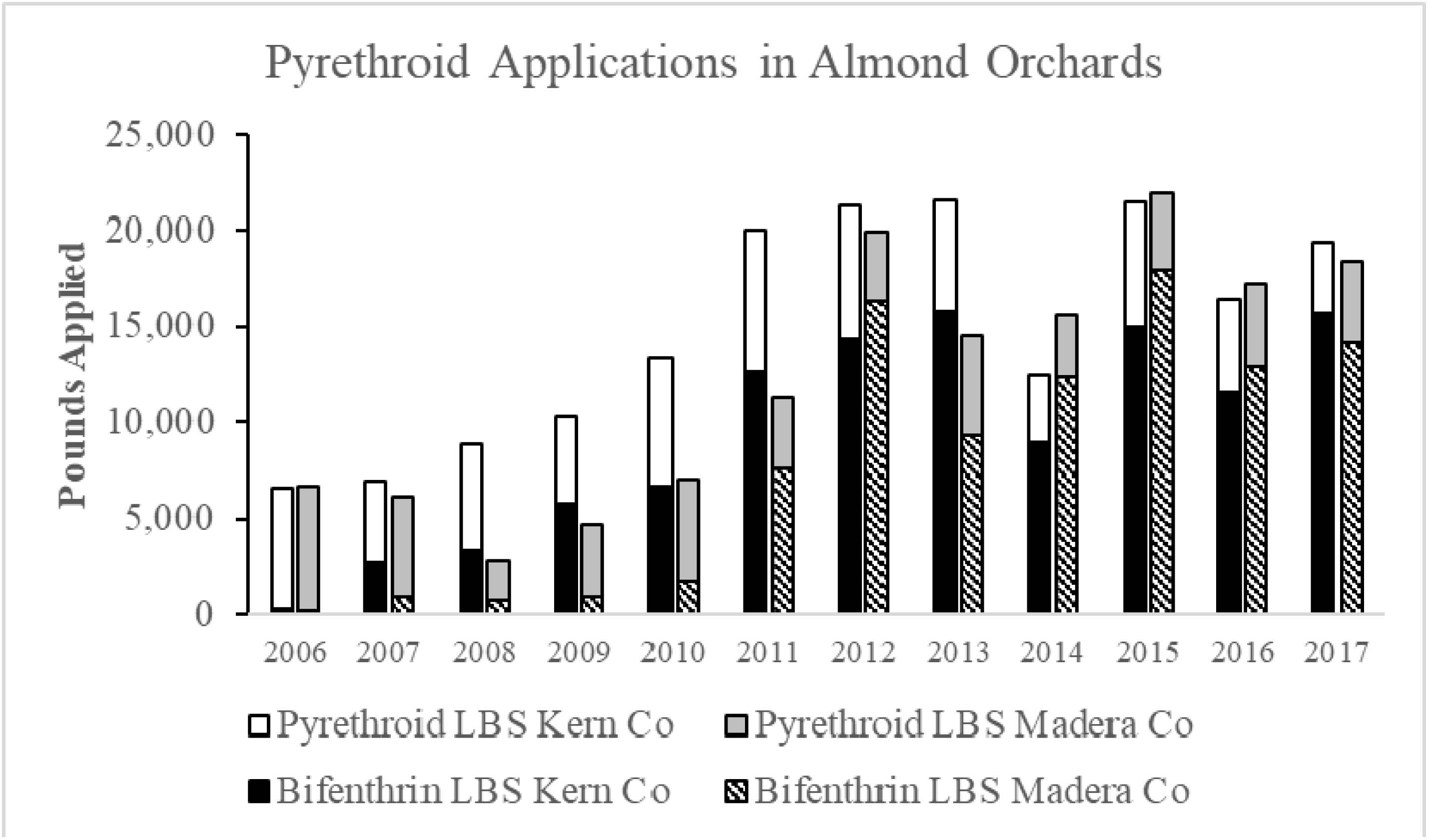
Bifenthrin use in Kern and Madera County almond orchards versus all other pyrethroids reported in the DPR pesticide use records (cyfluthrin, beta-cyfluthrin, (S)-cypermethrin, deltamethrin, esfenvalerate, fenpropathrin, lambda-cyhalothrin, gamma-cyhalothrin, permethrin) from 2006 to 2017.

## Discussion

A classic hallmark of positive selection is a reduction in nucleotide diversity in genome regions flanking genes that have been the targets of such selection (Kaplan et al., 1989). This shift in allele frequencies, or “selective sweep” (Kim and Stephan, 2002), is dependent on recombination rates in the genomic region as well as in the timing and strength of the underlying selective pressure. Classic selective sweeps or “hard sweeps” (where heterozygosity reaches near zero values) are rare and are not reproducible in experimental evolution studies, where for the most part soft sweeps and/or multi-locus resistance are obtained (Burke, 2012; Messer and Petrov, 2013). Examples of classic selective sweeps in insects are few. Hudson et al. (Hudson et al., 1994; Sáez et al., 2003), e.g., detected a ∼50-kb selective sweep surrounding the *SOD* locus in *Drosophila melanogaster*. Using high-density QTL markers, Lattorff et al. (Lattorff et al., 2015) detected a locus in the honeybee *Apis mellifera* where the allele frequency reached near fixation (allele frequency = 0.97) in a period of seven years under selection with the ectoparasitic mite *Varroa destructor*.

Within the context of pesticide selection, only a handful of studies have identified sweeps associated with resistance and, in all cases to date, the sweeps were relatively small (< 200 kb). Schlenke and Begun (Schlenke and Begun, 2004) reported a ∼100-kb selective sweep associated with a *Doc* transposable element inserted in *Cyp6g1* in *Drosophila simulans*, although the sweep was only marginally associated with DDT resistance. Soft, incomplete selective sweeps were found in the α-esterase gene cluster that contains the polymorphic *Lc*α*E7* gene encoding forms of the protein conferring organophosphate insecticide resistance in the Australian sheep blow fly *Lucilia cuprina* (Rose et al., 2011). A region containing the quantitative trait loci responsible for pyrethroid resistance in the malaria vector mosquito *Anopheles funestus* displayed characteristics of a selective sweep, which was then narrowed to the *CYP6P9A* and *CYP6P9B* clusters (Barnes et al., 2017). Song et al. (Song et al., 2015) analyzed nine Z chromosome-linked loci in different populations of the Old-World bollworm *Helicoverpa armigera* and detected a region possibly indicative of a selective sweep surrounding the Cyp303a1 locus.

In the navel orangeworm, a crop pest regularly subjected to repeated, widespread heavy use of a single insecticide, we have confirmed the existence of a large (∼1.3 Mb) region showing characteristics of a selective sweep in populations that show varying levels of resistance to bifenthrin. The sweep region contains 38 genes in the two susceptible lines sequenced and extends up to 43 genes in the resistant line. In this region, there are virtually no polymorphisms across 0.5 Mb (spanning 22 genes) among the three sequenced populations. The sweep then extends to both sides in the scaffold with increased – but still very low – polymorphism in ALM and FIG to the right of the hard sweep and increased polymorphism in the three populations to the left. In addition, the regions flanking both sides of the selective sweep show very high *F*_ST_ values between the resistant R347 and either the ALM or FIG populations. The genetic basis of the phenotypic differences (i.e., resistance to bifenthrin) very likely lies in part or entirely in this flanking region.

To evaluate the possible target or targets of selection that caused the observed patterns, we re-measured nucleotide diversity across each gene coding sequence and analyzed the amino acid sequences to verify the presence of non-silent mutations. The flanking region to the left of the hard-sweep, where the sweep is better characterized as a soft-sweep in all three populations, includes 15 genes, including cytochrome c oxidase, a cyclin-dependent kinase, a phosphomevalonate kinase and *Krüppel-like-9* transcription factor, among others. Of these, only cytochrome c oxidase has been directly associated with pyrethroid resistance in an insect (Pridgeon and Liu, 2003). The *Krüppel-like-9* transcription factor, however, has been implicated in regulation of P450 expression in mammals (Koh et al., 2014), and it showed segregating non-silent mutations that differ between the resistant and both susceptible strains. To the right of the sweep, where only R347 retains low nucleotide diversity, in addition to a gene encoding a small conductance calcium-activated potassium channel (SkCa2), there are two uncharacterized proteins and an extensin-like protein-coding gene. Although calcium and chloride ion channels have recently been found to interact with pyrethroids in mammals (Soderlund, 2012), to date there is no evidence that insect SkCa2 channels are targeted by pyrethroids.

The first gene in the hard-sweep region is *para* (paralytic), with the mutation that confers partial resistance to pyrethroids (*kdr*) in all three sequenced populations. The *kdr* locus has long been associated with resistance both to DDT and to other neurotoxic insecticides, including pyrethrins and pyrethroids (Kalra et al., 1967; Farnham, 1977), in many insects; *kdr* maps to a single recessive point mutation, resulting in an amino acid substitution from L to F in the S6 transmembrane segment of domain II of the *para* protein, which is a voltage-gated sodium channel, the main target of pyrethroids and DDT (Soderlund and Bloomquist, 1989). The *kdr* mutation is present in ALM, FIG, and R347 strains and absent in the reference genome and in the CPQ strain. Our bioassay results showed that resistance to DDT is associated with the presence of the *kdr* mutation (Table 3). However, differences in resistance to bifenthrin do not uniquely associated with the *kdr* mutation in the sequenced populations, indicating that additive factors are involved in resistance outcomes, including other beneficial mutations that might be responsible for the large selective sweep.

The *para* gene is followed by a cluster of three cytochrome P450 genes (*CYP6B53, CYP6B54* and *CYP6B56*), where *CYP6B56* has a π value of zero in R347. Although the precise substrate specificity of these P450s has not been defined, they belong to a CYP subfamily associated in other lepidopterans with pyrethroid metabolism (Li et al., 2004) and pyrethroid resistance (Zhang et al., 2010; Qiu et al., 2012). Assays with piperonyl butoxide, a P450 synergist, have implicated P450s in pyrethroid detoxification in *A. transitella* (Demkovich et al., 2015). Because there are no mutations in the coding sequences in these P450s in any of the sequenced strains, if their function has been the target of selection the causal variant may be located in a regulatory region. The fact that these three P450s are present as a tandem cluster indicates that they originated by duplication, an additional indication of their adaptive value.

The insecticidal properties of DDT were discovered in 1939, and it was heavily used by the military for vector control during World War II. Its release to civilian populations in 1945 led to widespread adoption for use against agricultural pests until 1972, when virtually all federal registrations for its use in the USA were cancelled by the Environmental Protection Agency. Despite its curtailed use, DDT and its derivatives remain in the environment, persisting in the soil as contaminants and moving from there across trophic levels and geographic regions (Mansouri et al., 2017). The *A. transitella* SPIRL-1966 line was established in the USDA laboratory facility in the San Joaquin Valley in 1966, during a period when the Central Valley area of California was subjected via crop-dusting to pesticides estimated at the time as amounting to “thousands of tons of DDT alone” (Cory et al., 1971).

Based on that information, and absent any information about recombination rates in *A. transitella*, we suggest that the large hard sweep documented in contemporary field populations of this insect is likely attributable to a stacking of selective forces that prevailed at the time of the founding of the SPIRL-1966 colony and that persisted under continuous selection until a new reinforcing selective pressure arrived in the form of pyrethroids. The fact that the genomic region of low variability in contemporary populations is nearly identical to the sequence in the SPIRL-1966 reference genome suggests that this reference strain had probably undergone a population bottleneck, possibly by selection with DDT, effectively reducing the number of alleles that were then maintained in the laboratory. Although SPIRL-1966 does not carry the *kdr* mutation, our findings are consistent with a scenario whereby it may have lost the mutation after generations of relaxed selection under laboratory conditions (Hanai et al., 2018).

High levels of resistance to pyrethroids in Kern County could have emerged as a result of a “perfect storm” of contributing factors—a newly cheap and widely available insecticide being used over an extensive area for a multivoltine pest requiring multiple applications for control in a high-value crop. Reports of selective sweeps of the size we have identified in this study are vanishingly rare. Tian et al. (Tian et al., 2009) reported a 1.1 Mb sweep in domesticated *Zea mays*, with similar evidence of multiple selection targets, the domestication of maize likely resulted from artificial selection exerted over thousands of years (Kistler et al., 2018), a significantly longer period than the time required for DDT and bifenthrin resistance to appear in *A. transitella* in Kern County. Within the context of pesticide selection, only a handful of studies have identified sweeps associated with resistance and, in most cases to date, the sweeps were relatively small (< 200 kb) (Schlenke and Begun, 2004) (Song et al., 2015; Barnes et al., 2017). In 2017, Kamdem et al. (Kamdem et al., 2017) compared urban and rural populations of mosquitoes in the *Anopheles gambiae* complex in Cameroon, where DDT is still used for malaria control, and identified a selective sweep containing circa 80 genes, among which was the *kdr* locus.

## Acknowledgements

We thank Dr. Mathew Hudson and Dr. Julian Catchen of the University of Illinois at Urbana-Champaign for helpful discussions and suggestions. We also thank Jeffrey Haas for helping with computing resources.

## Authors contributions

**BC, MD, MB and JS:** Wrote the manuscript; **BC, MD** and **JPGM** analyzed the data, **BC** and **MD** generated figures and graphical representations; **MD, JS, and KKOW** collected and processed samples and extracted DNA; **MB and HMR** conceived the study.

## Data Availability

The datasets generated for this study can be found in the NCBI sequence read archive (SRA), under accession numbers:

ALM: PRJNA544523, SAMN11842179, SRX5891168, SRR9117091;

FIG: PRJNA544523, SAMN11842181, SRX5891170, SRR9117089;

R347: PRJNA544523, SAMN11842180, SRX5891169, SRR9117090

Additional datasets analyzed in in this work can be accessed in the NCBI Bioproject accession: PRJNA548705

## Conflict of interests

The authors declare that they have no conflict of interests.

Mention of trade names and commercial products in this article is solely for the purpose of providing specific information and does not imply recommendation or endorsement by the US Department of Agriculture. US Department of Agriculture is an equal opportunity provider and employer

## Contribution to field statement

Using a population genomics approach, we have identified an unusually large selective sweep that spans several populations of the *A. transitella* genome. This sweep contains several genes with possible detoxification and target resistance functions. We suggest that this sweep is the result of previously selected regions undergoing a new round of selection with the recent heavy application of pyrethroids in the Central Valley of California. These findings provide evidence that decisions based in part on economic aspects of pest control can affect the genomic configuration of target and non-target organisms, possibly leading to a stacking of alleles that confer resistance and leading to the occurrence of extremely difficult to manage insects, mirroring the case of antibiotic resistance and the origin of *superbugs*. Our study also exemplifies the ability of humans to alter and accelerate the pace of evolutionary change in economically important insect species (Palumbi, 2001). Finally, this study illustrate a cost-effective method to assess population-level genomics without the need of re-sequencing single individuals independently.

