## Supplementary Material for "Selective sweeps in a nutshell; the genomic footprint of rapid insecticide resistance evolution in an insect"

**"Selective sweeps in a nutshell; the genomic footprint of rapid insecticide resistance evolution in a major insect pest of tree nuts"**

Bernarda Calla, Mark Demkovich, Joel P. Siegel, João Paulo Gomes Viana, Kim K.O. Walden, Hugh M. Robertson, May R. Berenbaum

Corresponding Author: Bernarda Calla

**List of Supplementary Materials:**
Tables S1-S8
Figures S1-S4

### Supplementary Tables

**Table S1.** Pooled DNA re-sequencing results.

| Strain | Library Name | Number of reads | Read length | Paired | Haploid genome length | Coverage |
|---|---|---|---|---|---|---|
| **Almond** | Almond_AAGGCCGTCA_L006_R1_001.fastq | 120,568,394 | 150 | 2 | 406,468,287 | 88.987 |
| **Fig** | Fig_AAGATCTGAG_L006_R1_001.fastq | 112,371,554 | 150 | 2 | 406,468,287 | 82.938 |
| **R347** | R347_AAGGTGCCTG_L006_R1_001.fastq | 119,586,826 | 150 | 2 | 406,468,287 | 88.263 |
| | **Total Reads (paired)** | **705,053,548** | | | | **260.1876594** |

**Table S2.** Nucleotide diversity across scaffold NW_013535362.1. The number of SNPs, read coverage and the calculated Tajima's π are shown for each of the sequenced populations.

| ID | gene_ID | Star | End | gene_name | ALM number of SNPS | ALM coverage | ALM_pi | FIG number of SNPS | FIG coverage | Fig_pi | R347 number of SNPS | R347 coverage | R347_pi |
|---|---|---|---|---|---|---|---|---|---|---|---|---|---|
| rna4429 | XM_013328369.1 | 3399243 | 3401314 | CYPB56 | 2 | 0.966 | 0.000119055 | 3 | 0.932 | 0.000146796 | 0 | 0.999 | 0 |
| rna4431 | XM_013328342.1 | 3413197 | 3420607 | protein ariadne-1%2C transcript variant X2 | 11 | 0.929 | 0.000169131 | 19 | 0.927 | 0.000340522 | 1 | 0.891 | 0.000012382 |
| rna4443 | XM_013328236.1 | 3549071 | 3550177 | protein kish-A | 0 | 0.536 | 0 | 5 | 0.921 | 0.00047616 | 1 | 1 | 0.000075361 |
| rna4430 | XM_013328341.1 | 3403909 | 3420607 | protein ariadne-1%2C transcript variant X1 | 19 | 0.851 | 0.000133218 | 43 | 0.92 | 0.000287535 | 13 | 0.865 | 0.000078983 |
| rna4427 | XM_013328250.1 | 3344559 | 3350148 | sodium channel protein para%2C transcript variant X2 | 1 | 0.873 | 0.000016742 | 7 | 0.958 | 0.000106802 | 6 | 0.838 | 0.000121451 |
| rna4426 | XM_013328249.1 | 3344559 | 3370779 | sodium channel protein para%2C transcript variant X1 | 33 | 0.829 | 0.000155447 | 41 | 0.898 | 0.000163303 | 30 | 0.826 | 0.000129155 |
| rna4444 | XM_013328238.1 | 3550487 | 3554188 | protein prenyltransferase alpha subunit repeat-containing protein 1 | 21 | 0.797 | 0.000809689 | 33 | 0.842 | 0.001342665 | 5 | 0.763 | 0.000158307 |
| rna4432 | XM_013328377.1 | 3421473 | 3423241 | uncharacterized LOC106129721 | 3 | 0.939 | 0.000147596 | 4 | 0.993 | 0.000186148 | 3 | 0.949 | 0.000168929 |
| rna4447 | XM_013328187.1 | 3745239 | 3796518 | protein unc-13 homolog A | 623 | 0.685 | 0.002149737 | 728 | 0.736 | 0.002379027 | 75 | 0.764 | 0.000174787 |
| rna4448 | XM_013328228.1 | 3800374 | 3817393 | actin-binding Rho-activating protein-like | 214 | 0.665 | 0.002477321 | 304 | 0.707 | 0.003822464 | 27 | 0.729 | 0.000196256 |
| rna4433 | XM_013328378.1 | 3423369 | 3430390 | isoleucine--tRNA ligase%2C cytoplasmic | 11 | 0.683 | 0.000210947 | 32 | 0.746 | 0.000671049 | 15 | 0.719 | 0.000265044 |
| rna4445 | XM_013328171.1 | 3584697 | 3630468 | uncharacterized LOC106129575 | 485 | 0.659 | 0.001909073 | 652 | 0.653 | 0.002842798 | 97 | 0.675 | 0.000319536 |
| rna4434 | XM_013328269.1 | 3433176 | 3489393 | dmX-like protein 2 | 408 | 0.637 | 0.001241575 | 449 | 0.667 | 0.001385853 | 121 | 0.646 | 0.000323764 |
| rna4442 | XM_013328235.1 | 3529722 | 3548710 | ras GTPase-activating protein 1 | 173 | 0.693 | 0.001636741 | 241 | 0.761 | 0.00197034 | 52 | 0.741 | 0.00035081 |
| rna4428 | XM_013328169.1 | 3387986 | 3396394 | CYPB54-55 | 31 | 0.877 | 0.000374048 | 67 | 0.945 | 0.000832382 | 30 | 0.889 | 0.000397261 |
| rna4449 | XM_013328172.1 | 3826528 | 3849030 | gonadotropin-releasing hormone II receptor-like | 326 | 0.757 | 0.002598439 | 460 | 0.75 | 0.004254889 | 87 | 0.778 | 0.00047073 |
| rna4438 | XM_013328231.1 | 3490744 | 3524106 | endophilin-A%2C transcript variant X1 | 288 | 0.792 | 0.001364922 | 273 | 0.854 | 0.001120561 | 106 | 0.847 | 0.000511522 |
| rna4440 | XM_013328232.1 | 3490744 | 3524106 | endophilin-A%2C transcript variant X2 | 288 | 0.792 | 0.001364922 | 273 | 0.854 | 0.001120561 | 106 | 0.847 | 0.000511522 |
| rna4439 | XM_013328233.1 | 3490744 | 3524106 | endophilin-A%2C transcript variant X3 | 288 | 0.792 | 0.001364922 | 273 | 0.854 | 0.001120561 | 106 | 0.847 | 0.000511522 |
| rna4446 | XM_013328188.1 | 3631722 | 3638019 | uncharacterized LOC106129590 | 57 | 0.629 | 0.001609691 | 75 | 0.677 | 0.002214825 | 24 | 0.611 | 0.000589032 |

| | | | | | | | | | | | | | |
|---|---|---|---|---|---|---|---|---|---|---|---|---|---|
| rna4455 | XM_013328217.1 | 4270710 | 4272681 | tyrosine--tRNA ligase%2C cytoplasmic | 15 | 0.589 | 0.002191836 | 22 | 0.915 | 0.002546646 | 8 | 0.902 | 0.000683768 |
| rna4390 | XM_013328351.1 | 2580309 | 2581660 | UDP-GlcNAc:betaGal beta-1%2C3-N-acetylglucosaminyltransferase-like protein 1 | 7 | 0.949 | 0.001699828 | 5 | 0.919 | 0.00140442 | 6 | 0.865 | 0.001191301 |
| rna4418 | XM_013328423.1 | 3159383 | 3164900 | uncharacterized LOC106129757 | 26 | 0.673 | 0.001504173 | 21 | 0.711 | 0.001399366 | 25 | 0.73 | 0.001236382 |
| rna4337 | XM_013328277.1 | 1505319 | 1507681 | conserved oligomeric Golgi complex subunit 5 | 24 | 0.647 | 0.003325352 | 36 | 0.924 | 0.004487335 | 12 | 0.89 | 0.001648673 |
| rna4425 | XM_013328229.1 | 3304902 | 3307354 | phosphomevalonate kinase | 28 | 0.914 | 0.002209508 | 28 | 0.967 | 0.002526449 | 17 | 0.712 | 0.001704432 |
| rna4413 | XM_013328280.1 | 3097871 | 3099727 | GPI mannosyltransferase 2%2C transcript variant X2 | 10 | 0.893 | 0.001245343 | 7 | 0.907 | 0.000848715 | 6 | 0.912 | 0.001815043 |
| rna4414 | XM_013328279.1 | 3097905 | 3099727 | GPI mannosyltransferase 2%2C transcript variant X1 | 9 | 0.891 | 0.001221095 | 7 | 0.906 | 0.000866193 | 6 | 0.919 | 0.001834536 |
| rna4441 | XM_013328234.1 | 3525167 | 3528547 | polyubiquitin-C | 51 | 0.639 | 0.002994905 | 40 | 0.676 | 0.002750336 | 26 | 0.67 | 0.002126739 |
| rna4391 | XM_013328352.1 | 2581733 | 2585900 | stress-induced-phosphoprotein 1-like | 50 | 0.893 | 0.00302288 | 54 | 0.942 | 0.003067504 | 31 | 0.899 | 0.002209818 |
| rna4422 | XM_013328326.1 | 3204704 | 3279423 | protein groucho%2C transcript variant X1 | 512 | 0.615 | 0.00255649 | 570 | 0.629 | 0.002639457 | 492 | 0.618 | 0.00259059 |
| rna4451 | XM_013328392.1 | 4090436 | 4108229 | uncharacterized LOC106129735 | 333 | 0.696 | 0.007056351 | 396 | 0.76 | 0.006815791 | 213 | 0.745 | 0.002636165 |
| rna4423 | XM_013328328.1 | 3204704 | 3258508 | protein groucho%2C transcript variant X3 | 410 | 0.604 | 0.002993113 | 461 | 0.614 | 0.003024779 | 386 | 0.605 | 0.002909226 |
| rna4424 | XM_013328327.1 | 3204704 | 3256842 | protein groucho%2C transcript variant X2 | 393 | 0.593 | 0.003028286 | 443 | 0.603 | 0.003050611 | 369 | 0.593 | 0.002923422 |
| rna4383 | XM_013328409.1 | 2419495 | 2425584 | putative elongator complex protein 1%2C transcript variant X1 | 90 | 0.895 | 0.004315263 | 94 | 0.923 | 0.004142576 | 66 | 0.893 | 0.003205471 |
| rna4384 | XM_013328410.1 | 2419495 | 2425584 | putative elongator complex protein 1%2C transcript variant X2 | 90 | 0.895 | 0.004315263 | 94 | 0.923 | 0.004142576 | 66 | 0.893 | 0.003205471 |
| rna4421 | XM_013328247.1 | 3197609 | 3199024 | Krueppel-like factor 9 | 24 | 0.833 | 0.005024417 | 23 | 0.898 | 0.004206684 | 23 | 0.965 | 0.003498993 |
| rna4450 | XM_013328189.1 | 3852565 | 4034421 | small conductance calcium-activated potassium channel protein | 4722 | 0.628 | 0.008780796 | 5092 | 0.629 | 0.00959187 | 2798 | 0.693 | 0.003576904 |
| rna4408 | XM_013328167.1 | 3044389 | 3048122 | uncharacterized LOC106129572 | 45 | 0.801 | 0.002517566 | 27 | 0.887 | 0.001446331 | 36 | 0.948 | 0.003757452 |
| rna4453 | XM_013328173.1 | 4247890 | 4251407 | uncharacterized protein DDB_G0286591-like | 71 | 0.792 | 0.005590469 | 75 | 0.86 | 0.005145974 | 42 | 0.849 | 0.003790941 |
| rna4530 | XM_013328414.1 | 5881767 | 5883523 | DTW domain-containing protein 2 | 25 | 0.789 | 0.004663524 | 25 | 0.873 | 0.004628095 | 20 | 0.701 | 0.004146948 |
| rna4508 | XM_013328386.1 | 5483318 | 5487648 | odorant receptor 4-like | 138 | 0.779 | 0.00653974 | 120 | 0.773 | 0.006490545 | 83 | 0.822 | 0.004175595 |
| rna4331 | XM_013328225.1 | 1473705 | 1476159 | m7GpppN-mRNA hydrolase | 29 | 0.927 | 0.003160619 | 27 | 0.933 | 0.003546826 | 21 | 0.901 | 0.004217887 |
| rna4501 | XM_013328240.1 | 5313693 | 5315051 | mitochondrial inner membrane protein OXA1L | 25 | 0.993 | 0.005148463 | 17 | 0.722 | 0.004908737 | 15 | 0.712 | 0.004220994 |

| | | | | | | | | | | | | | |
|---|---|---|---|---|---|---|---|---|---|---|---|---|---|
| rna4517 | XM_013328185.1 | 5724518 | 5727850 | uncharacterized LOC106129587 | 34 | 0.743 | 0.00451717 | 54 | 0.939 | 0.004745916 | 41 | 0.852 | 0.004393735 |
| rna4405 | XM_013328243.1 | 3002489 | 3003978 | cytochrome c oxidase assembly protein COX16 homolog%2C mitochondrial | 27 | 0.712 | 0.006641446 | 19 | 0.83 | 0.005655548 | 23 | 0.85 | 0.004474807 |
| rna4456 | XM_013328325.1 | 4275218 | 4323747 | RNA polymerase II elongation factor ELL | 966 | 0.783 | 0.006161608 | 978 | 0.829 | 0.005349858 | 571 | 0.798 | 0.004598154 |
| rna4396 | XM_013328380.1 | 2593141 | 2603646 | myelin expression factor 2-like%2C transcript variant X2 | 145 | 0.608 | 0.007917822 | 152 | 0.588 | 0.008019857 | 110 | 0.601 | 0.004679833 |
| rna4321 | XM_013328346.1 | 1399170 | 1408734 | uncharacterized LOC106129698 | 161 | 0.71 | 0.006730182 | 160 | 0.689 | 0.006518281 | 85 | 0.622 | 0.004759801 |
| rna4330 | XM_013328241.1 | 1471195 | 1473165 | syntaxin-18 | 30 | 0.758 | 0.007262027 | 33 | 0.697 | 0.008038343 | 18 | 0.688 | 0.004929163 |
| rna4420 | XM_013328361.1 | 3164948 | 3193977 | unconventional myosin-IXa-like%2C transcript variant X1 | 579 | 0.756 | 0.005938819 | 548 | 0.797 | 0.005037003 | 496 | 0.774 | 0.005021211 |
| rna4419 | XM_013328362.1 | 3164948 | 3193977 | unconventional myosin-IXa-like%2C transcript variant X2 | 579 | 0.756 | 0.005938819 | 548 | 0.797 | 0.005037003 | 496 | 0.774 | 0.005021211 |
| rna4338 | XM_013328278.1 | 1508705 | 1518258 | GTPase HRas | 131 | 0.659 | 0.006153678 | 126 | 0.693 | 0.006000635 | 100 | 0.672 | 0.005151254 |
| rna4340 | XM_013328422.1 | 1519937 | 1520766 | dolichol-phosphate mannosyltransferase subunit 3%2C transcript variant X2 | 17 | 0.882 | 0.007364632 | 18 | 0.861 | 0.007774272 | 12 | 0.972 | 0.005275164 |
| rna4454 | XM_013328174.1 | 4268658 | 4270209 | uncharacterized LOC106129578 | 43 | 0.922 | 0.007847321 | 45 | 0.918 | 0.0065733 | 41 | 0.852 | 0.005321223 |
| rna4536 | XM_013328244.1 | 6012409 | 6013623 | uncharacterized protein C19orf52 | 20 | 0.737 | 0.004995216 | 31 | 0.891 | 0.007180479 | 19 | 0.792 | 0.005331318 |
| rna4521 | XM_013328393.1 | 5745189 | 5751175 | gbkey=mRNA | 85 | 0.62 | 0.005618301 | 85 | 0.623 | 0.005251222 | 94 | 0.693 | 0.005670913 |
| rna4393 | XM_013328382.1 | 2593138 | 2598806 | myelin expression factor 2-like%2C transcript variant X4 | 79 | 0.626 | 0.008517212 | 81 | 0.592 | 0.008954112 | 65 | 0.617 | 0.005697184 |
| rna4298 | XM_013328239.1 | 880578 | 883070 | protein bud22 | 49 | 0.868 | 0.005870367 | 49 | 0.893 | 0.005227974 | 44 | 0.878 | 0.005884998 |
| rna4339 | XM_013328421.1 | 1519661 | 1520744 | dolichol-phosphate mannosyltransferase subunit 3%2C transcript variant X1 | 24 | 0.885 | 0.008233695 | 25 | 0.869 | 0.008556691 | 16 | 0.956 | 0.006077829 |
| rna4378 | XM_013328165.1 | 2399919 | 2400599 | uncharacterized LOC106129569 | 13 | 0.921 | 0.007158843 | 15 | 0.938 | 0.007078859 | 11 | 0.927 | 0.006090894 |
| rna4496 | XM_013328333.1 | 5271861 | 5273537 | neugrin | 32 | 0.769 | 0.0075507 | 40 | 0.908 | 0.007070333 | 30 | 0.882 | 0.006332797 |
| rna4380 | XM_013328417.1 | 2410837 | 2412344 | beta-ureidopropionase-like | 25 | 0.854 | 0.005200978 | 29 | 0.936 | 0.005376323 | 23 | 0.885 | 0.006505778 |
| rna4381 | XM_013328418.1 | 2412449 | 2413979 | eukaryotic translation initiation factor 2 subunit 2 | 19 | 0.673 | 0.004709099 | 25 | 0.943 | 0.004881273 | 23 | 0.94 | 0.006689415 |
| rna4406 | XM_013328367.1 | 3012239 | 3045757 | tyrosine-protein kinase CSK%2C transcript variant X1 | 558 | 0.648 | 0.007588042 | 530 | 0.702 | 0.006464824 | 463 | 0.68 | 0.006805918 |
| rna4385 | XM_013328242.1 | 2425702 | 2429091 | leucine-rich repeat protein 1 | 68 | 0.792 | 0.007598856 | 67 | 0.804 | 0.006949772 | 53 | 0.814 | 0.006885488 |
| rna4392 | XM_013328399.1 | 2587577 | 2592737 | integrator complex subunit 3 homolog | 84 | 0.716 | 0.008476183 | 90 | 0.864 | 0.007165859 | 88 | 0.719 | 0.007058129 |
| rna4511 | XM_013328182.1 | 5504764 | 5505982 | lipase 1-like | 17 | 0.587 | 0.007667123 | 31 | 0.974 | 0.009696404 | 27 | 0.996 | 0.007102324 |

| | | | | | | | | | | | | | |
|---|---|---|---|---|---|---|---|---|---|---|---|---|---|
| rna4359 | XM_013328358.1 | 1827009 | 1830838 | vacuolar protein sorting-associated protein 33B%2C transcript variant X2 | 73 | 0.552 | 0.01019639 | 60 | 0.46 | 0.011606181 | 48 | 0.539 | 0.007282337 |
| rna4360 | XM_013328357.1 | 1827979 | 1830838 | vacuolar protein sorting-associated protein 33B%2C transcript variant X1 | 73 | 0.74 | 0.01019639 | 60 | 0.616 | 0.011606181 | 48 | 0.722 | 0.007282337 |
| rna4300 | XM_013328258.1 | 887424 | 889772 | ribose-phosphate pyrophosphokinase 2 | 42 | 0.755 | 0.007613161 | 43 | 0.858 | 0.006280155 | 43 | 0.883 | 0.007461008 |
| rna4348 | XM_013328291.1 | 1577783 | 1581428 | BTB/POZ domain-containing protein 2-like%2C transcript variant X1 | 81 | 0.68 | 0.011976623 | 81 | 0.681 | 0.011489997 | 64 | 0.672 | 0.00752791 |
| rna4407 | XM_013328368.1 | 3019509 | 3045757 | tyrosine-protein kinase CSK%2C transcript variant X2 | 492 | 0.681 | 0.008041054 | 472 | 0.742 | 0.006853115 | 415 | 0.723 | 0.007566126 |
| rna4452 | XM_013328198.1 | 4240437 | 4246388 | extensin-like | 260 | 0.789 | 0.015132831 | 260 | 0.809 | 0.013739108 | 178 | 0.84 | 0.007591015 |
| rna4389 | XM_013328350.1 | 2572396 | 2580080 | gbkey=mRNA | 148 | 0.667 | 0.00859795 | 151 | 0.665 | 0.008366142 | 130 | 0.68 | 0.007648406 |
| rna4354 | XM_013328201.1 | 1651101 | 1653941 | alpha-(1%2C3)-fucosyltransferase C-like | 98 | 0.778 | 0.012263638 | 106 | 0.841 | 0.013055453 | 63 | 0.659 | 0.007756923 |
| rna4534 | XM_013328246.1 | 6000856 | 6002196 | uncharacterized LOC106129643 | 27 | 0.961 | 0.00609803 | 26 | 0.936 | 0.005062659 | 27 | 0.919 | 0.00785931 |
| rna4493 | XM_013328336.1 | 5236222 | 5238866 | uncharacterized LOC106129692 | 67 | 0.783 | 0.008596993 | 71 | 0.952 | 0.008128525 | 65 | 0.846 | 0.007874551 |
| rna4525 | XM_013328271.1 | 5775698 | 5788089 | uncharacterized LOC106129662%2C transcript variant X2 | 196 | 0.592 | 0.007924001 | 204 | 0.625 | 0.007401777 | 207 | 0.574 | 0.0080822 |
| rna4526 | XM_013328270.1 | 5775698 | 5786474 | uncharacterized LOC106129662%2C transcript variant X1 | 196 | 0.681 | 0.007924001 | 204 | 0.719 | 0.007403688 | 207 | 0.659 | 0.0080822 |
| rna4500 | XM_013328224.1 | 5310573 | 5313063 | U3 small nucleolar RNA-associated protein 15 homolog | 52 | 0.679 | 0.009316192 | 75 | 0.933 | 0.008723168 | 45 | 0.729 | 0.00821631 |
| rna4522 | XM_013328200.1 | 5753445 | 5759187 | uncharacterized LOC106129599 | 106 | 0.669 | 0.008081293 | 79 | 0.596 | 0.006874992 | 82 | 0.542 | 0.008634491 |
| rna4281 | XM_013328149.1 | 70277 | 73054 | uncharacterized LOC106129551 | 70 | 0.949 | 0.006904584 | 76 | 0.979 | 0.00770778 | 78 | 0.967 | 0.00865995 |
| rna4347 | XM_013328292.1 | 1575910 | 1581428 | BTB/POZ domain-containing protein 2-like%2C transcript variant X2 | 116 | 0.576 | 0.012664871 | 119 | 0.583 | 0.012238872 | 96 | 0.565 | 0.009015599 |
| rna4507 | XM_013328387.1 | 5479669 | 5483229 | pancreatic lipase-related protein 2-like | 112 | 0.858 | 0.010547971 | 129 | 0.906 | 0.011299822 | 95 | 0.929 | 0.009177413 |
| rna4386 | XM_013328214.1 | 2429317 | 2434186 | G kinase-anchoring protein 1-like | 132 | 0.752 | 0.012244815 | 128 | 0.73 | 0.011291087 | 95 | 0.724 | 0.009191826 |
| rna4328 | XM_013328196.1 | 1466142 | 1467679 | RNA-binding protein NOB1 | 35 | 0.888 | 0.009440154 | 36 | 0.916 | 0.009234366 | 30 | 0.845 | 0.009248617 |
| rna4343 | XM_013328379.1 | 1553769 | 1557284 | membrane-bound alkaline phosphatase-like | 74 | 0.617 | 0.008592208 | 76 | 0.608 | 0.009571366 | 57 | 0.588 | 0.009338058 |
| rna4403 | XM_013328219.1 | 2977372 | 2991119 | ADAM 17-like protease | 242 | 0.457 | 0.011319361 | 219 | 0.5 | 0.008955323 | 195 | 0.512 | 0.009365391 |
| rna4345 | XM_013328290.1 | 1563041 | 1565652 | membrane-bound alkaline phosphatase-like | 66 | 0.873 | 0.009084757 | 62 | 0.892 | 0.007516861 | 55 | 0.755 | 0.010169592 |

| | | | | | | | | | | | | | |
|---|---|---|---|---|---|---|---|---|---|---|---|---|---|
| rna4409 | XM_013328206.1 | 3049409 | 3053897 | DET1 homolog | 92 | 0.804 | 0.0053506 | 79 | 0.875 | 0.003573386 | 82 | 0.766 | 0.010415605 |
| rna4382 | XM_013328215.1 | 2415442 | 2418993 | 28S ribosomal protein S30%2C mitochondrial | 129 | 0.852 | 0.013025224 | 125 | 0.84 | 0.012615374 | 99 | 0.787 | 0.010437139 |
| rna4531 | XM_013328415.1 | 5884495 | 5886643 | 40S ribosomal protein S15 | 66 | 0.752 | 0.011841779 | 74 | 0.872 | 0.010363965 | 61 | 0.874 | 0.010457711 |
| rna4387 | XM_013328412.1 | 2436696 | 2532139 | A disintegrin and metalloproteinase with thrombospondin motifs 7-like | 2049 | 0.597 | 0.011803109 | 2047 | 0.617 | 0.011169004 | 1781 | 0.604 | 0.010484045 |
| rna4529 | XM_013328186.1 | 5830408 | 5878979 | uncharacterized LOC106129588 | 1115 | 0.587 | 0.010991996 | 1164 | 0.586 | 0.011132651 | 1045 | 0.581 | 0.011062135 |
| rna4467 | XM_013328376.1 | 4842481 | 4850978 | dual specificity mitogen-activated protein kinase kinase dSOR1 | 286 | 0.812 | 0.01048265 | 292 | 0.873 | 0.009530437 | 234 | 0.817 | 0.011341106 |
| rna4352 | XM_013328364.1 | 1632464 | 1635170 | putative nuclease HARBI1 | 75 | 0.743 | 0.01204676 | 91 | 0.762 | 0.012629692 | 65 | 0.685 | 0.011566315 |
| rna4400 | XM_013328253.1 | 2883596 | 2887227 | uncharacterized LOC106129648 | 135 | 0.766 | 0.015635148 | 137 | 0.811 | 0.01543218 | 102 | 0.792 | 0.011600085 |
| rna4329 | XM_013328197.1 | 1467658 | 1470732 | UPF0528 protein CG10038 | 93 | 0.84 | 0.014087576 | 109 | 0.801 | 0.014739893 | 57 | 0.712 | 0.011617053 |
| rna4461 | XM_013328287.1 | 4473947 | 4534011 | uncharacterized LOC106129672 | 2056 | 0.697 | 0.013389586 | 2151 | 0.729 | 0.012438962 | 1599 | 0.701 | 0.01164602 |
| rna4509 | XM_013328180.1 | 5490242 | 5493926 | odorant receptor 4-like | 163 | 0.842 | 0.015328105 | 149 | 0.796 | 0.014657463 | 135 | 0.875 | 0.011770148 |
| rna4460 | XM_013328288.1 | 4419275 | 4437090 | protein giant-lens | 483 | 0.573 | 0.013003857 | 487 | 0.576 | 0.012881567 | 421 | 0.594 | 0.012005888 |
| rna4416 | XM_013328281.1 | 3101631 | 3107700 | methyl-CpG-binding domain protein 3%2C transcript variant X1 | 141 | 0.739 | 0.006844267 | 119 | 0.823 | 0.005241431 | 121 | 0.717 | 0.012030669 |
| rna4415 | XM_013328282.1 | 3101631 | 3107700 | methyl-CpG-binding domain protein 3%2C transcript variant X2 | 141 | 0.739 | 0.006844267 | 119 | 0.823 | 0.005241431 | 121 | 0.717 | 0.012030669 |
| rna4361 | XM_013328345.1 | 1831183 | 1847961 | probable isocitrate dehydrogenase [NAD] subunit alpha%2C mitochondrial%2C transcript variant X3 | 432 | 0.581 | 0.01368344 | 454 | 0.619 | 0.013274785 | 387 | 0.614 | 0.012033377 |
| rna4411 | XM_013328168.1 | 3067370 | 3078948 | tubulin polyglutamylase TTLL13-like | 262 | 0.693 | 0.006962775 | 211 | 0.671 | 0.00557638 | 260 | 0.689 | 0.012051446 |
| rna4362 | XM_013328343.1 | 1831299 | 1847961 | probable isocitrate dehydrogenase [NAD] subunit alpha%2C mitochondrial%2C transcript variant X1 | 430 | 0.579 | 0.013766682 | 453 | 0.617 | 0.013375931 | 385 | 0.611 | 0.012077437 |
| rna4506 | XM_013328203.1 | 5471655 | 5477166 | pancreatic lipase-related protein 2-like | 123 | 0.432 | 0.016173196 | 137 | 0.444 | 0.017888798 | 111 | 0.46 | 0.012177467 |
| rna4342 | XM_013328158.1 | 1549021 | 1552044 | membrane-bound alkaline phosphatase-like | 95 | 0.765 | 0.011345937 | 99 | 0.908 | 0.009984343 | 97 | 0.943 | 0.012457168 |
| rna4510 | XM_013328181.1 | 5500322 | 5503424 | putative odorant receptor 92a | 191 | 0.872 | 0.023127907 | 184 | 0.869 | 0.021780068 | 144 | 0.864 | 0.012824788 |

| | | | | | | | | | | | | | |
|---|---|---|---|---|---|---|---|---|---|---|---|---|---|
| rna4333 | XM_013328427.1 | 1476505 | 1484339 | glycosylated lysosomal membrane protein B-like%2C transcript variant X2 | 200 | 0.704 | 0.010096267 | 221 | 0.727 | 0.011198945 | 181 | 0.648 | 0.012865083 |
| rna4332 | XM_013328426.1 | 1476505 | 1484348 | glycosylated lysosomal membrane protein B-like%2C transcript variant X1 | 201 | 0.704 | 0.010105206 | 221 | 0.727 | 0.011187163 | 182 | 0.648 | 0.012878185 |
| rna4350 | XM_013328300.1 | 1621486 | 1629660 | monocarboxylate transporter 1-like%2C transcript variant X1 | 256 | 0.681 | 0.015368042 | 244 | 0.645 | 0.015738436 | 227 | 0.68 | 0.013095432 |
| rna4537 | XM_013328428.1 | 6014289 | 6025097 | exception=annotated by transcript or proteomic data | 315 | 0.696 | 0.011987582 | 321 | 0.726 | 0.011711206 | 328 | 0.71 | 0.013096788 |
| rna4322 | XM_013328354.1 | 1415367 | 1429459 | zinc finger protein 135-like%2C transcript variant X2 | 453 | 0.594 | 0.018622603 | 476 | 0.59 | 0.018443142 | 267 | 0.517 | 0.013110547 |
| rna4357 | XM_013328161.1 | 1795791 | 1810707 | uncharacterized LOC106129566 | 526 | 0.66 | 0.018768043 | 519 | 0.7 | 0.017989018 | 408 | 0.622 | 0.013126036 |
| rna4476 | XM_013328309.1 | 5164520 | 5213764 | casein kinase I isoform gamma-3%2C transcript variant X6 | 1778 | 0.689 | 0.01438096 | 1813 | 0.728 | 0.014184591 | 1446 | 0.729 | 0.01322272 |
| rna4477 | XM_013328313.1 | 5164520 | 5213764 | casein kinase I isoform gamma-3%2C transcript variant X9 | 1778 | 0.689 | 0.01438096 | 1813 | 0.728 | 0.014184591 | 1446 | 0.729 | 0.01322272 |
| rna4482 | XM_013328304.1 | 5164520 | 5213751 | casein kinase I isoform gamma-3%2C transcript variant X2 | 1778 | 0.689 | 0.014386049 | 1813 | 0.728 | 0.014189739 | 1446 | 0.729 | 0.013227508 |
| rna4481 | XM_013328305.1 | 5164520 | 5213751 | casein kinase I isoform gamma-3%2C transcript variant X3 | 1778 | 0.689 | 0.014386049 | 1813 | 0.728 | 0.014189739 | 1446 | 0.729 | 0.013227508 |
| rna4480 | XM_013328310.1 | 5164520 | 5213751 | casein kinase I isoform gamma-3%2C transcript variant X7 | 1778 | 0.689 | 0.014386049 | 1813 | 0.728 | 0.014189739 | 1446 | 0.729 | 0.013227508 |
| rna4479 | XM_013328311.1 | 5164520 | 5213751 | casein kinase I isoform gamma-3%2C transcript variant X8 | 1778 | 0.689 | 0.014386049 | 1813 | 0.728 | 0.014189739 | 1446 | 0.729 | 0.013227508 |
| rna4478 | XM_013328314.1 | 5164520 | 5213751 | casein kinase I isoform gamma-3%2C transcript variant X10 | 1778 | 0.689 | 0.014386049 | 1813 | 0.728 | 0.014189739 | 1446 | 0.729 | 0.013227508 |
| rna4485 | XM_013328317.1 | 5164520 | 5213751 | casein kinase I isoform gamma-3%2C transcript variant X13 | 1778 | 0.689 | 0.014386049 | 1813 | 0.728 | 0.014189739 | 1446 | 0.729 | 0.013227508 |
| rna4486 | XM_013328318.1 | 5164520 | 5213751 | casein kinase I isoform gamma-3%2C transcript variant X14 | 1778 | 0.689 | 0.014386049 | 1813 | 0.728 | 0.014189739 | 1446 | 0.729 | 0.013227508 |
| rna4484 | XM_013328319.1 | 5164520 | 5213751 | casein kinase I isoform gamma-3%2C transcript variant X15 | 1778 | 0.689 | 0.014386049 | 1813 | 0.728 | 0.014189739 | 1446 | 0.729 | 0.013227508 |
| rna4483 | XM_013328320.1 | 5164520 | 5213751 | casein kinase I isoform gamma-3%2C transcript variant X16 | 1778 | 0.689 | 0.014386049 | 1813 | 0.728 | 0.014189739 | 1446 | 0.729 | 0.013227508 |
| rna4487 | XM_013328303.1 | 5164520 | 5213739 | casein kinase I isoform gamma-3%2C transcript variant X1 | 1778 | 0.689 | 0.014391142 | 1813 | 0.728 | 0.014194494 | 1446 | 0.729 | 0.013231931 |

| | | | | | | | | | | | | | |
|---|---|---|---|---|---|---|---|---|---|---|---|---|---|
| rna4488 | XM_013328308.1 | 5164520 | 5213731 | casein kinase I isoform gamma-3%2C transcript variant X5 | 1778 | 0.689 | 0.014394538 | 1813 | 0.727 | 0.014197666 | 1446 | 0.729 | 0.013234881 |
| rna4363 | XM_013328344.1 | 1831299 | 1842059 | probable isocitrate dehydrogenase [NAD] subunit alpha%2C mitochondrial%2C transcript variant X2 | 354 | 0.66 | 0.01500007 | 366 | 0.705 | 0.014674234 | 315 | 0.7 | 0.013238679 |
| rna4489 | XM_013328315.1 | 5164520 | 5213712 | casein kinase I isoform gamma-3%2C transcript variant X11 | 1778 | 0.689 | 0.014402613 | 1813 | 0.727 | 0.014205205 | 1446 | 0.729 | 0.013241893 |
| rna4490 | XM_013328307.1 | 5164520 | 5213697 | casein kinase I isoform gamma-3%2C transcript variant X4 | 1778 | 0.688 | 0.014408993 | 1813 | 0.727 | 0.014211163 | 1446 | 0.729 | 0.013247434 |
| rna4491 | XM_013328316.1 | 5164520 | 5213680 | casein kinase I isoform gamma-3%2C transcript variant X12 | 1778 | 0.688 | 0.014416232 | 1813 | 0.727 | 0.014217921 | 1446 | 0.729 | 0.013253719 |
| rna4351 | XM_013328301.1 | 1621486 | 1629516 | monocarboxylate transporter 1-like%2C transcript variant X2 | 253 | 0.677 | 0.015556926 | 240 | 0.64 | 0.015965943 | 224 | 0.676 | 0.013263462 |
| rna4397 | XM_013328166.1 | 2628373 | 2634398 | putative odorant receptor 85d | 171 | 0.636 | 0.014448566 | 180 | 0.667 | 0.014342764 | 151 | 0.625 | 0.013279379 |
| rna4402 | XM_013328221.1 | 2922570 | 2976883 | dynein heavy chain%2C cytoplasmic | 1839 | 0.608 | 0.015209379 | 1734 | 0.624 | 0.013681591 | 1319 | 0.61 | 0.013491432 |
| rna4417 | XM_013328283.1 | 3128643 | 3157699 | phospholipid-transporting ATPase ID | 550 | 0.61 | 0.006533429 | 479 | 0.63 | 0.005536496 | 591 | 0.605 | 0.013728037 |
| rna4367 | XM_013328394.1 | 1871706 | 1877740 | uncharacterized family 31 glucosidase KIAA1161-like%2C transcript variant X1 | 227 | 0.771 | 0.014574838 | 206 | 0.719 | 0.01482246 | 173 | 0.669 | 0.013877593 |
| rna4366 | XM_013328395.1 | 1871706 | 1878974 | uncharacterized family 31 glucosidase KIAA1161-like%2C transcript variant X2 | 227 | 0.64 | 0.014574838 | 206 | 0.597 | 0.01482246 | 173 | 0.556 | 0.013877593 |
| rna4369 | XM_013328416.1 | 1888912 | 1894807 | uncharacterized family 31 glucosidase KIAA1161-like | 187 | 0.646 | 0.015901968 | 190 | 0.675 | 0.01622541 | 165 | 0.618 | 0.013886772 |
| rna4516 | XM_013328385.1 | 5717817 | 5721577 | 40S ribosomal protein S6 | 141 | 0.808 | 0.016680076 | 126 | 0.747 | 0.015159258 | 119 | 0.771 | 0.013921093 |
| rna4457 | XM_013328363.1 | 4325443 | 4337300 | arginine-glutamic acid dipeptide repeats protein-like | 404 | 0.648 | 0.01402339 | 374 | 0.668 | 0.011080246 | 346 | 0.711 | 0.013923976 |
| rna4466 | XM_013328178.1 | 4834313 | 4841636 | chromatin assembly factor 1 subunit A | 268 | 0.691 | 0.014858696 | 302 | 0.781 | 0.01333414 | 254 | 0.765 | 0.013975477 |
| rna4323 | XM_013328353.1 | 1417895 | 1429459 | zinc finger protein 135-like%2C transcript variant X1 | 434 | 0.639 | 0.020277646 | 455 | 0.625 | 0.020355006 | 255 | 0.553 | 0.014106421 |
| rna4320 | XM_013328400.1 | 1384695 | 1394057 | A-kinase anchor protein 17A%2C transcript variant X1 | 253 | 0.593 | 0.013673796 | 230 | 0.567 | 0.014024146 | 220 | 0.588 | 0.014323947 |
| rna4319 | XM_013328401.1 | 1384695 | 1394057 | A-kinase anchor protein 17A%2C transcript variant X2 | 253 | 0.593 | 0.013673796 | 230 | 0.567 | 0.014024146 | 220 | 0.588 | 0.014323947 |

| | | | | | | | | | | | | | |
|---|---|---|---|---|---|---|---|---|---|---|---|---|---|
| rna4318 | XM_013328402.1 | 1384695 | 1394057 | A-kinase anchor protein 17A%2C transcript variant X3 | 253 | 0.593 | 0.013673796 | 230 | 0.567 | 0.014024146 | 220 | 0.588 | 0.014323947 |
| rna4317 | XM_013328403.1 | 1384695 | 1394057 | A-kinase anchor protein 17A%2C transcript variant X4 | 253 | 0.593 | 0.013673796 | 230 | 0.567 | 0.014024146 | 220 | 0.588 | 0.014323947 |
| rna4316 | XM_013328404.1 | 1384695 | 1394057 | A-kinase anchor protein 17A%2C transcript variant X5 | 253 | 0.593 | 0.013673796 | 230 | 0.567 | 0.014024146 | 220 | 0.588 | 0.014323947 |
| rna4515 | XM_013328424.1 | 5705067 | 5712195 | geranylgeranyl transferase type-1 subunit beta%2C transcript variant X1 | 319 | 0.777 | 0.019859861 | 354 | 0.801 | 0.020228957 | 302 | 0.816 | 0.014330747 |
| rna4514 | XM_013328425.1 | 5705067 | 5712195 | geranylgeranyl transferase type-1 subunit beta%2C transcript variant X2 | 319 | 0.777 | 0.019859861 | 354 | 0.801 | 0.020228957 | 302 | 0.816 | 0.014330747 |
| rna4341 | XM_013328157.1 | 1524364 | 1540086 | integral membrane protein DGCR2/IDD-like | 500 | 0.627 | 0.016272457 | 506 | 0.645 | 0.015666339 | 380 | 0.617 | 0.014409885 |
| rna4305 | XM_013328265.1 | 917570 | 930179 | uncharacterized LOC106129656 | 383 | 0.516 | 0.017309033 | 347 | 0.477 | 0.017272682 | 344 | 0.577 | 0.014794716 |
| rna4356 | XM_013328227.1 | 1667064 | 1770730 | gonadotropin-releasing hormone II receptor-like | 2501 | 0.502 | 0.013139661 | 2641 | 0.517 | 0.013387471 | 2322 | 0.517 | 0.014889395 |
| rna4326 | XM_013328321.1 | 1449157 | 1465233 | probable ATP-dependent RNA helicase YTHDC2 | 395 | 0.492 | 0.017093656 | 402 | 0.486 | 0.016996883 | 324 | 0.51 | 0.015137028 |
| rna4518 | XM_013328199.1 | 5731508 | 5738487 | ubiquitin-like modifier-activating enzyme ATG7 | 341 | 0.744 | 0.018723624 | 391 | 0.784 | 0.020415932 | 323 | 0.786 | 0.015780398 |
| rna4399 | XM_013328195.1 | 2740022 | 2773970 | hemicentin-2-like | 1411 | 0.7 | 0.018305714 | 1269 | 0.711 | 0.01579168 | 1105 | 0.681 | 0.015959163 |
| rna4388 | XM_013328405.1 | 2562109 | 2571267 | regulator of nonsense transcripts 1 homolog | 260 | 0.623 | 0.01448283 | 234 | 0.63 | 0.011783985 | 212 | 0.593 | 0.016005798 |
| rna4375 | XM_013328207.1 | 2213901 | 2250952 | cell adhesion molecule 3-like | 1117 | 0.553 | 0.017373042 | 1189 | 0.582 | 0.017199421 | 1037 | 0.565 | 0.016310751 |
| rna4335 | XM_013328275.1 | 1490555 | 1505353 | lon protease homolog%2C mitochondrial%2C transcript variant X1 | 441 | 0.495 | 0.019835248 | 460 | 0.506 | 0.018761867 | 353 | 0.514 | 0.016329611 |
| rna4336 | XM_013328276.1 | 1490555 | 1505353 | lon protease homolog%2C mitochondrial%2C transcript variant X2 | 441 | 0.495 | 0.019835248 | 460 | 0.506 | 0.018761867 | 353 | 0.514 | 0.016329611 |
| rna4527 | XM_013328272.1 | 5788149 | 5817440 | synaptic vesicle glycoprotein 2C-like%2C transcript variant X1 | 1090 | 0.654 | 0.016902493 | 1121 | 0.652 | 0.01678627 | 1069 | 0.663 | 0.016341617 |
| rna4364 | XM_013328430.1 | 1849499 | 1858817 | nucleolar protein 6 | 387 | 0.631 | 0.01841051 | 373 | 0.676 | 0.017427999 | 315 | 0.666 | 0.016504625 |
| rna4519 | XM_013328211.1 | 5739062 | 5743761 | 39S ribosomal protein L2%2C mitochondrial%2C transcript variant X2 | 144 | 0.41 | 0.023212282 | 155 | 0.473 | 0.023363934 | 123 | 0.462 | 0.01668886 |
| rna4462 | XM_013328175.1 | 4535269 | 4543692 | elongation factor Tu GTP-binding domain-containing protein 1 | 296 | 0.568 | 0.017343452 | 274 | 0.57 | 0.015638323 | 301 | 0.665 | 0.016740826 |
| rna4346 | XM_013328159.1 | 1568083 | 1573750 | katanin p60 ATPase-containing subunit A-like 2 | 253 | 0.786 | 0.017294694 | 244 | 0.793 | 0.015920924 | 229 | 0.799 | 0.016830798 |

| | | | | | | | | | | | | | |
|---|---|---|---|---|---|---|---|---|---|---|---|---|---|
| rna4520 | XM_013328210.1 | 5739062 | 5742744 | 39S ribosomal protein L2%2C mitochondrial%2C transcript variant X1 | 138 | 0.493 | 0.02372714 | 148 | 0.568 | 0.023889944 | 119 | 0.568 | 0.016839515 |
| rna4465 | XM_013328384.1 | 4817352 | 4831584 | uncharacterized LOC106129727 | 595 | 0.638 | 0.018827495 | 612 | 0.672 | 0.017066858 | 481 | 0.619 | 0.016952938 |
| rna4311 | XM_013328154.1 | 1063427 | 1075258 | uncharacterized LOC106129557 | 549 | 0.639 | 0.021962126 | 500 | 0.634 | 0.020626919 | 461 | 0.699 | 0.016967 |
| rna4474 | XM_013328347.1 | 5144998 | 5163317 | uncharacterized LOC106129699%2C transcript variant X1 | 851 | 0.696 | 0.018764671 | 789 | 0.691 | 0.017536698 | 694 | 0.709 | 0.017073215 |
| rna4475 | XM_013328349.1 | 5145472 | 5163317 | uncharacterized LOC106129699%2C transcript variant X2 | 839 | 0.688 | 0.019198705 | 776 | 0.686 | 0.017860861 | 685 | 0.701 | 0.017474922 |
| rna4289 | XM_013328208.1 | 507747 | 509855 | uncharacterized LOC106129607 | 108 | 0.786 | 0.019687015 | 90 | 0.794 | 0.017459236 | 108 | 0.843 | 0.017548592 |
| rna4463 | XM_013328176.1 | 4664744 | 4714572 | hemicentin-2-like | 2152 | 0.606 | 0.020292481 | 2118 | 0.62 | 0.018615133 | 1829 | 0.63 | 0.017549343 |
| rna4458 | XM_013328284.1 | 4354138 | 4400298 | uncharacterized protein YHR080C-like%2C transcript variant X1 | 1614 | 0.56 | 0.016665457 | 1534 | 0.571 | 0.013892726 | 1408 | 0.586 | 0.017572023 |
| rna4358 | XM_013328356.1 | 1811917 | 1826990 | phosphatidylinositol 3-kinase catalytic subunit type 3 | 608 | 0.589 | 0.022799511 | 628 | 0.608 | 0.022059835 | 474 | 0.564 | 0.017755397 |
| rna4470 | XM_013328294.1 | 4961729 | 5141211 | furin-like protease 1%2C isoform 1-CRR%2C transcript variant X1 | 6652 | 0.56 | 0.018331297 | 6785 | 0.572 | 0.018259894 | 5726 | 0.587 | 0.01777704 |
| rna4471 | XM_013328295.1 | 4961729 | 5141211 | furin-like protease 1%2C isoform 1-CRR%2C transcript variant X2 | 6652 | 0.56 | 0.018331297 | 6785 | 0.572 | 0.018259894 | 5726 | 0.587 | 0.01777704 |
| rna4468 | XM_013328297.1 | 4961729 | 5141211 | furin-like protease 1%2C isoform 1-CRR%2C transcript variant X4 | 6652 | 0.56 | 0.018331297 | 6785 | 0.572 | 0.018259894 | 5726 | 0.587 | 0.01777704 |
| rna4469 | XM_013328299.1 | 4961729 | 5141211 | furin-like protease 1%2C isoform 1-CRR%2C transcript variant X6 | 6652 | 0.56 | 0.018331297 | 6785 | 0.572 | 0.018259894 | 5726 | 0.587 | 0.01777704 |
| rna4353 | XM_013328365.1 | 1635267 | 1650031 | AP-3 complex subunit beta-2 | 497 | 0.482 | 0.019673635 | 517 | 0.493 | 0.019788226 | 408 | 0.484 | 0.01778626 |
| rna4473 | XM_013328296.1 | 4961729 | 5139591 | furin-like protease 1%2C isoform 1-CRR%2C transcript variant X3 | 6605 | 0.558 | 0.018454357 | 6735 | 0.57 | 0.018383854 | 5691 | 0.585 | 0.017901082 |
| rna4472 | XM_013328298.1 | 4961729 | 5139591 | furin-like protease 1%2C isoform 1-CRR%2C transcript variant X5 | 6605 | 0.558 | 0.018454357 | 6735 | 0.57 | 0.018383854 | 5691 | 0.585 | 0.017901082 |
| rna4334 | XM_013328156.1 | 1485721 | 1490391 | probable tRNA (guanine(26)-N(2))-dimethyltransferase | 215 | 0.821 | 0.018499936 | 223 | 0.84 | 0.017830596 | 172 | 0.79 | 0.017903263 |
| rna4459 | XM_013328286.1 | 4367881 | 4400298 | uncharacterized protein YHR080C-like%2C transcript variant X2 | 1022 | 0.514 | 0.01637634 | 992 | 0.521 | 0.014090233 | 924 | 0.538 | 0.017914521 |
| rna4379 | XM_013328205.1 | 2401400 | 2408494 | microfibrillar-associated protein 1 | 217 | 0.598 | 0.01490628 | 209 | 0.613 | 0.013053112 | 195 | 0.579 | 0.017974082 |

| | | | | | | | | | | | | | |
|---|---|---|---|---|---|---|---|---|---|---|---|---|---|
| rna4528 | XM_013328274.1 | 5788149 | 5807240 | synaptic vesicle glycoprotein 2C-like%2C transcript variant X2 | 781 | 0.667 | 0.018624867 | 796 | 0.654 | 0.018646787 | 777 | 0.679 | 0.017990336 |
| rna4494 | XM_013328340.1 | 5239658 | 5249675 | alpha-tocopherol transfer protein-like | 543 | 0.667 | 0.024193356 | 535 | 0.671 | 0.023512712 | 475 | 0.728 | 0.017992505 |
| rna4377 | XM_013328202.1 | 2397004 | 2399474 | acyl-CoA Delta(11) desaturase-like | 95 | 0.807 | 0.013121725 | 91 | 0.826 | 0.012078733 | 105 | 0.853 | 0.018038584 |
| rna4370 | XM_013328371.1 | 1915714 | 1997644 | gonadotropin-releasing hormone receptor%2C transcript variant X1 | 3020 | 0.565 | 0.017658915 | 3207 | 0.582 | 0.018459627 | 2565 | 0.538 | 0.018231373 |
| rna4371 | XM_013328372.1 | 1915714 | 1997644 | gonadotropin-releasing hormone receptor%2C transcript variant X2 | 3020 | 0.565 | 0.017658915 | 3207 | 0.582 | 0.018459627 | 2565 | 0.538 | 0.018231373 |
| rna4297 | XM_013328212.1 | 878042 | 880103 | 60S ribosomal protein L36 | 74 | 0.586 | 0.017021303 | 66 | 0.605 | 0.014981928 | 98 | 0.756 | 0.018501056 |
| rna4398 | XM_013328251.1 | 2650908 | 2658501 | odorant receptor 46a%2C isoform A-like | 374 | 0.816 | 0.017949239 | 348 | 0.817 | 0.015522419 | 310 | 0.798 | 0.018573135 |
| rna4513 | XM_013328183.1 | 5584108 | 5587497 | zinc metalloproteinase nas-26-like | 170 | 0.727 | 0.020259458 | 191 | 0.693 | 0.022580203 | 156 | 0.725 | 0.018650785 |
| rna4313 | XM_013328155.1 | 1323121 | 1327462 | uncharacterized LOC106129558 | 271 | 0.757 | 0.027620403 | 255 | 0.767 | 0.02655026 | 160 | 0.666 | 0.018674776 |
| rna4283 | XM_013328150.1 | 117071 | 202199 | COUP transcription factor 2 | 3901 | 0.639 | 0.020747332 | 3839 | 0.644 | 0.020308331 | 3463 | 0.656 | 0.018676525 |
| rna4524 | XM_013328407.1 | 5764182 | 5774900 | ejaculatory bulb-specific protein 3-like%2C transcript variant X1 | 424 | 0.663 | 0.017350391 | 448 | 0.695 | 0.016681601 | 457 | 0.695 | 0.018691693 |
| rna4349 | XM_013328160.1 | 1589746 | 1617720 | ras-specific guanine nucleotide-releasing factor 2-like | 1042 | 0.63 | 0.018533524 | 1086 | 0.664 | 0.016560191 | 874 | 0.629 | 0.018795288 |
| rna4299 | XM_013328256.1 | 886270 | 900020 | histone H4 transcription factor%2C transcript variant X1 | 577 | 0.604 | 0.020228768 | 579 | 0.659 | 0.018517783 | 538 | 0.662 | 0.018815606 |
| rna4344 | XM_013328396.1 | 1559700 | 1562822 | membrane-bound alkaline phosphatase-like | 138 | 0.786 | 0.021208463 | 136 | 0.814 | 0.019345777 | 129 | 0.795 | 0.019003645 |
| rna4523 | XM_013328408.1 | 5763005 | 5774900 | ejaculatory bulb-specific protein 3-like%2C transcript variant X2 | 482 | 0.667 | 0.017765032 | 494 | 0.693 | 0.016863614 | 521 | 0.702 | 0.019232355 |
| rna4324 | XM_013328355.1 | 1429485 | 1433468 | D-amino-acid oxidase | 180 | 0.769 | 0.022401047 | 171 | 0.748 | 0.021670132 | 150 | 0.721 | 0.019578476 |
| rna4512 | XM_013328245.1 | 5523782 | 5567906 | lachesin-like | 1999 | 0.614 | 0.022743925 | 2034 | 0.622 | 0.022294619 | 1683 | 0.619 | 0.019783072 |
| rna4533 | XM_013328302.1 | 5936832 | 5986228 | tyrosine-protein phosphatase non-receptor type 9-like | 1915 | 0.593 | 0.018338658 | 1984 | 0.597 | 0.019156148 | 1753 | 0.582 | 0.020455713 |
| rna4365 | XM_013328213.1 | 1859120 | 1867969 | uncharacterized family 31 glucosidase KIAA1161-like | 333 | 0.553 | 0.01865628 | 301 | 0.504 | 0.018463486 | 278 | 0.574 | 0.020574957 |
| rna4325 | XM_013328323.1 | 1434200 | 1447798 | TFIIH basal transcription factor complex helicase XPD subunit | 470 | 0.483 | 0.023278871 | 455 | 0.464 | 0.023244102 | 403 | 0.505 | 0.020644034 |
| rna4312 | XM_013328390.1 | 1276242 | 1283154 | collagenase-like | 281 | 0.542 | 0.023178141 | 278 | 0.537 | 0.023880987 | 249 | 0.578 | 0.020653982 |

| | | | | | | | | | | | | | |
|---|---|---|---|---|---|---|---|---|---|---|---|---|---|
| rna4355 | XM_013328209.1 | 1658562 | 1661332 | gbkey=mRNA | 132 | 0.851 | 0.016475081 | 118 | 0.766 | 0.016984606 | 109 | 0.725 | 0.02076845 |
| rna4368 | XM_013328190.1 | 1881341 | 1887617 | uncharacterized family 31 glucosidase KIAA1161-like | 271 | 0.578 | 0.019633061 | 263 | 0.578 | 0.019645999 | 252 | 0.593 | 0.020841519 |
| rna4314 | XM_013328230.1 | 1353928 | 1366053 | cytochrome P450 6B2-like | 506 | 0.545 | 0.025493476 | 436 | 0.532 | 0.021572619 | 386 | 0.52 | 0.020849574 |
| rna4307 | XM_013328263.1 | 932579 | 937549 | survival motor neuron protein%2C transcript variant X1 | 268 | 0.717 | 0.023064884 | 286 | 0.677 | 0.024100881 | 210 | 0.621 | 0.021044792 |
| rna4306 | XM_013328264.1 | 932579 | 937549 | survival motor neuron protein%2C transcript variant X2 | 268 | 0.717 | 0.023064884 | 286 | 0.677 | 0.024100881 | 210 | 0.621 | 0.021044792 |
| rna4308 | XM_013328413.1 | 938212 | 946286 | uncharacterized LOC106129749 | 448 | 0.713 | 0.023860812 | 489 | 0.762 | 0.024650573 | 347 | 0.713 | 0.021213124 |
| rna4495 | XM_013328337.1 | 5249756 | 5270691 | alpha-tocopherol transfer protein-like | 1070 | 0.64 | 0.023430361 | 1089 | 0.665 | 0.021801526 | 932 | 0.655 | 0.021302354 |
| rna4532 | XM_013328391.1 | 5891606 | 5936552 | bumetanide-sensitive sodium-(potassium)-chloride cotransporter | 1427 | 0.474 | 0.021341498 | 1417 | 0.459 | 0.021724603 | 1381 | 0.472 | 0.021385654 |
| rna4499 | XM_013328329.1 | 5273552 | 5310024 | DENN domain-containing protein 4C%2C transcript variant X1 | 1542 | 0.563 | 0.022557767 | 1455 | 0.543 | 0.021318098 | 1466 | 0.579 | 0.0219706 |
| rna4497 | XM_013328330.1 | 5273552 | 5310024 | DENN domain-containing protein 4C%2C transcript variant X2 | 1542 | 0.563 | 0.022557767 | 1455 | 0.543 | 0.021318098 | 1466 | 0.579 | 0.0219706 |
| rna4498 | XM_013328332.1 | 5273552 | 5310024 | DENN domain-containing protein 4C%2C transcript variant X3 | 1542 | 0.563 | 0.022557767 | 1455 | 0.543 | 0.021318098 | 1466 | 0.579 | 0.0219706 |
| rna4401 | XM_013328254.1 | 2892093 | 2895881 | uncharacterized LOC106129649 | 177 | 0.741 | 0.018112111 | 173 | 0.774 | 0.01880833 | 148 | 0.73 | 0.02242318 |
| rna4286 | XM_013328255.1 | 296812 | 298122 | uncharacterized LOC106129650 | 75 | 0.73 | 0.024917858 | 71 | 0.683 | 0.026239023 | 60 | 0.686 | 0.023446898 |
| rna4292 | XM_013328420.1 | 782839 | 797582 | organic cation transporter protein-like | 940 | 0.76 | 0.024639982 | 874 | 0.727 | 0.024891206 | 791 | 0.746 | 0.023580162 |
| rna4295 | XM_013328191.1 | 851105 | 875577 | organic cation transporter protein-like%2C transcript variant X1 | 1225 | 0.591 | 0.024989271 | 1111 | 0.58 | 0.023653437 | 1056 | 0.607 | 0.023840622 |
| rna4296 | XM_013328194.1 | 852790 | 875577 | organic cation transporter protein-like%2C transcript variant X2 | 1131 | 0.579 | 0.024994831 | 1019 | 0.567 | 0.023516426 | 975 | 0.595 | 0.023936719 |
| rna4301 | XM_013328257.1 | 892898 | 900020 | histone H4 transcription factor%2C transcript variant X2 | 437 | 0.702 | 0.026294065 | 427 | 0.694 | 0.026057796 | 405 | 0.716 | 0.024658554 |
| rna4464 | XM_013328383.1 | 4810659 | 4814482 | peptidyl-prolyl cis-trans isomerase G | 204 | 0.703 | 0.022109041 | 215 | 0.754 | 0.021423466 | 147 | 0.554 | 0.024671476 |
| rna4504 | XM_013328334.1 | 5452362 | 5457029 | pancreatic lipase-related protein 2-like | 314 | 0.79 | 0.023964422 | 329 | 0.781 | 0.024215162 | 275 | 0.771 | 0.025397708 |
| rna4502 | XM_013328397.1 | 5438263 | 5441330 | pancreatic lipase-related protein 2-like | 178 | 0.691 | 0.028531789 | 175 | 0.694 | 0.02718466 | 160 | 0.666 | 0.025959637 |

| | | | | | | | | | | | | | |
|---|---|---|---|---|---|---|---|---|---|---|---|---|---|
| rna4293 | XM_013328374.1 | 808877 | 830079 | organic cation transporter protein-like%2C transcript variant X1 | 1004 | 0.551 | 0.026870029 | 967 | 0.537 | 0.025994914 | 886 | 0.537 | 0.026360269 |
| rna4291 | XM_013328431.1 | 633693 | 760199 | lachesin-like | 5027 | 0.438 | 0.026772375 | 4845 | 0.431 | 0.026513013 | 4484 | 0.433 | 0.026460228 |
| rna4503 | XM_013328179.1 | 5446800 | 5451715 | pancreatic lipase-related protein 2-like | 299 | 0.699 | 0.030131834 | 314 | 0.727 | 0.03010721 | 296 | 0.736 | 0.026774886 |
| rna4287 | XM_013328252.1 | 328410 | 329562 | uncharacterized LOC106129647 | 69 | 0.635 | 0.030412102 | 66 | 0.547 | 0.03066087 | 67 | 0.714 | 0.026841506 |
| rna4294 | XM_013328375.1 | 808877 | 828008 | organic cation transporter protein-like%2C transcript variant X2 | 855 | 0.518 | 0.027457545 | 839 | 0.508 | 0.02640907 | 759 | 0.5 | 0.026858788 |
| rna4280 | XM_013328223.1 | 37998 | 40521 | proteasome subunit beta type-3 | 166 | 0.697 | 0.030671404 | 145 | 0.657 | 0.028231876 | 161 | 0.736 | 0.027159099 |
| rna4290 | XM_013328153.1 | 518291 | 555617 | limbic system-associated membrane protein-like | 2167 | 0.628 | 0.027881285 | 2052 | 0.605 | 0.02781822 | 2100 | 0.633 | 0.027433434 |
| rna4410 | XM_013328222.1 | 3055117 | 3067241 | cyclin-dependent kinase 7 | 123 | 0.462 | 0.004064722 | 90 | 0.451 | 0.003204948 | 122 | 0.388 | na |
| rna4278 | XM_013328389.1 | 2546 | 14627 | gbkey=mRNA | 119 | 0.179 | na | 95 | 0.153 | na | 161 | 0.285 | na |
| rna4284 | XM_013328151.1 | 264305 | 268869 | uncharacterized LOC106129553 | 48 | 0.207 | na | 54 | 0.212 | na | 74 | 0.229 | na |
| rna4285 | XM_013328289.1 | 269265 | 295932 | mediator of DNA damage checkpoint protein 1-like | 645 | 0.347 | na | 596 | 0.355 | na | 635 | 0.396 | na |
| rna4288 | XM_013328152.1 | 348091 | 349107 | uncharacterized LOC106129555 | 0 | 0 | na | 0 | 0 | na | 0 | 0 | na |
| rna4304 | XM_013328259.1 | 900124 | 912396 | spermine oxidase-like%2C transcript variant X1 | 208 | 0.223 | na | 191 | 0.207 | na | 199 | 0.229 | na |
| rna4302 | XM_013328261.1 | 900124 | 920223 | spermine oxidase-like%2C transcript variant X2 | 522 | 0.388 | na | 487 | 0.371 | na | 442 | 0.396 | na |
| rna4303 | XM_013328262.1 | 900124 | 913399 | spermine oxidase-like%2C transcript variant X3 | 208 | 0.206 | na | 191 | 0.191 | na | 199 | 0.212 | na |
| rna4309 | XM_013328218.1 | 949659 | 955857 | uncharacterized LOC106129616 | 25 | 0.042 | na | 21 | 0.058 | na | 26 | 0.061 | na |
| rna4315 | XM_013328406.1 | 1367764 | 1382026 | chymotrypsin BII-like | 288 | 0.246 | na | 237 | 0.219 | na | 198 | 0.219 | na |
| rna4327 | XM_013328324.1 | 1462726 | 1464554 | uncharacterized LOC106129685 | 3 | 0.04 | na | 5 | 0.021 | na | 0 | 0 | na |
| rna4373 | XM_013328373.1 | 1953448 | 1954299 | uncharacterized LOC106129718 | 0 | 0 | na | 0 | 0 | na | 0 | 0 | na |
| rna4374 | XM_013328162.1 | 2097656 | 2098856 | uncharacterized LOC106129567 | 13 | 0.06 | na | 13 | 0.062 | na | 14 | 0.062 | na |
| rna4376 | XM_013328164.1 | 2257330 | 2262771 | uncharacterized LOC106129568 | 12 | 0.071 | na | 13 | 0.064 | na | 7 | 0.044 | na |
| rna4404 | XM_013328220.1 | 2996162 | 2999913 | polyubiquitin-B | 58 | 0.353 | na | 56 | 0.37 | na | 49 | 0.369 | na |
| rna4412 | XM_013328359.1 | 3079983 | 3096923 | ubiquilin-1-like | 132 | 0.243 | na | 133 | 0.259 | na | 156 | 0.245 | na |
| rna4435 | XM_013328268.1 | 3476653 | 3481076 | uncharacterized LOC106129658 | 0 | 0.009 | na | 0 | 0.004 | na | 0 | 0.009 | na |

| | | | | | | | | | | | | | |
|---|---|---|---|---|---|---|---|---|---|---|---|---|---|
| rna4437 | XM_013328266.1 | 3479317 | 3481542 | uncharacterized LOC106129657%2C transcript variant X1 | 0 | 0 | na | 0 | 0 | na | 0 | 0 | na |
| rna4436 | XM_013328267.1 | 3479317 | 3481542 | uncharacterized LOC106129657%2C transcript variant X2 | 0 | 0 | na | 0 | 0 | na | 0 | 0 | na |
| rna4492 | XM_013328338.1 | 5215501 | 5234703 | alpha-tocopherol transfer protein-like | 558 | 0.365 | na | 593 | 0.387 | na | 400 | 0.356 | na |
| rna4505 | XM_013328335.1 | 5457125 | 5461303 | pancreatic lipase-related protein 2-like | 91 | 0.351 | na | 93 | 0.343 | na | 84 | 0.369 | na |

**Table S3.** Scanning of single point mutations relative to reference genome or major allele in the hard sweep region in scaffold NW_013535362.1 in *A. transitella*

| Scaffold | start | end | strand | functional_annotation(NCBI) | Transcript_ID | point mutations or deletions in coding regions |
|---|---|---|---|---|---|---|
| NW_013535362.1 | 3164948 | 3193977 | - | unconventional myosin-IXa-like | XM_013328361.1 | polymorphisms end here, R is identical to SPIRL at both ends. |
| NW_013535362.1 | 3197609 | 3199024 | - | Krueppel-like factor 9 | XM_013328247.1 | 3 non-silent point mutations |
| NW_013535362.1 | 3204704 | 3279423 | - | protein groucho | XM_013328326.1 | ------- |
| NW_013535362.1 | 3304902 | 3307354 | + | phosphomevalonate kinase | XM_013328229.1 | E127N (conservative mutation) |
| NW_013535362.1 | 3344559 | 3370779 | + | sodium channel protein para | XM_013328249.1 | L934F (kdr mutation) |
| NW_013535362.1 | 3387986 | 3396394 | - | CYP6B54 and CYP6B55 | XM_013328169.1 | none |
| NW_013535362.1 | 3399243 | 3401314 | - | CYP6B56 | XM_013328369.1 | none |
| NW_013535362.1 | 3403909 | 3420607 | - | protein ariadne-1 | XM_013328341.1 | none |
| NW_013535362.1 | 3421473 | 3423241 | - | uncharacterized LOC106129721 | XM_013328377.1 | none |
| NW_013535362.1 | 3423369 | 3430390 | + | isoleucine--tRNA ligase | XM_013328378.1 | none |
| NW_013535362.1 | 3433176 | 3489393 | - | dmX-like protein 2 | XM_013328269.1 | none |
| NW_013535362.1 | 3476653 | 3481076 | + | uncharacterized LOC106129658 | XM_013328268.1 | none |
| NW_013535362.1 | 3479317 | 3481542 | - | uncharacterized LOC106129657 | XM_013328266.1 | none |
| NW_013535362.1 | 3490744 | 3524106 | + | endophilin-A | XM_013328231.1 | none |
| NW_013535362.1 | 3525167 | 3528547 | - | polyubiquitin-C | XM_013328234.1 | Q13H (conservative mutation) |
| NW_013535362.1 | 3529722 | 3548710 | + | ras GTPase-activating protein 1 | XM_013328235.1 | none |
| NW_013535362.1 | 3549071 | 3550177 | - | protein kish-A | XM_013328236.1 | none |
| NW_013535362.1 | 3550487 | 3554188 | + | prenyltransferase alpha subunit | XM_013328238.1 | none |
| NW_013535362.1 | 3584697 | 3630468 | + | uncharacterized LOC106129575 | XM_013328171.1 | 1 alternative start codon (ATG->GTC segregating in 20% of ALM and 20% FIG) |
| NW_013535362.1 | 3631722 | 3638019 | - | uncharacterized LOC106129590 | XM_013328188.1 | none |
| NW_013535362.1 | 3745239 | 3796518 | + | protein unc-13 homolog A | XM_013328187.1 | none |
| NW_013535362.1 | 3800374 | 3817393 | + | actin-binding Rho-activating protein | XM_013328228.1 | none |
| NW_013535362.1 | 3826528 | 3849030 | - | gonadotropin-releasing hormone II | XM_013328172.1 | none |
| NW_013535362.1 | 3852565 | 4034421 | - | small conductance calcium-activate | XM_013328189.1 | all silent mutations |

**Table S4.** Twenty most differentiated SNPs between the reference genome (SPIRL-1966), ALM, FIG and R using Mahalanobis distances (PCAdapt, 1.1, Luu et al., 2017). **pos**: position along scaffold, **rc**: reference character, **allele count:** number of alleles found in all populations; **allele states**: allele characters in all populations (sorted by counts in all populations); **del sum**: sum of deletions in all populations (should be zero, if not the postion may not be reliable); **snp type**: SNP type: "pop" a SNP within or between the populations; "rc" a SNP between the reference sequence character and the consensus of at least one populaton; "rc|pop" both; **major alleles**: most frequent allele in all populations [ALM FIG R]; **minor alleles**: second most frequent allele in all populations [ALM FIG R]

| Scaffold# | pos | Ref | allele count | allele states | SNP type | major alleles (maa) | minor alleles (mia) | FDR pval. | gene model functional annotation |
|---|---|---|---|---|---|---|---|---|---|
| NW_013535379.1 | 775604 | A | 3 | A/C/G | pop | AAA | CCC | 0.01859 | inter-genic, nearby CYP341 cluster of P450s |
| NW_013535334.1 | 84304 | G | 3 | G/T/A | pop | GGG | TTT | 0.01970 | zinc finger 224-like |
| NW_013535359.1 | 304369 | A | 2 | A/T | pop | AAA | TTT | 0.01970 | inter-genic |
| NW_013535509.1 | 95606 | C | 2 | C/A | rc\|pop | ACC | CAA | 0.01970 | inter-genic just upstream of Ecdisteroid Kinase cluster |
| NW_013535530.1 | 170651 | T | 2 | T/C | pop | TTT | CCC | 0.01970 | inter-genic |
| NW_013535492.1 | 2659949 | T | 3 | A/G/T | rc\|pop | AAA | TGG | 0.03339 | intronic mannose-1-phosphate guanyltransferase alpha |
| NW_013535509.1 | 185180 | T | 2 | T/A | rc\|pop | TTA | AAT | 0.03339 | intergenic in Ecdisyteroid kinase gene cluster |
| NW_013535854.1 | 148606 | C | 2 | C/T | pop | CCC | TTT | 0.03339 | intergenic, just downstream of sorting nexin-2 |
| NW_013535700.1 | 310754 | A | 2 | A/G | pop | AAA | GGG | 0.04845 | intergenic, just downstream of protein king tubby |
| NW_013535323.1 | 5604262 | T | 2 | T/G | pop | TTT | GGG | 0.05023 | intergenic |
| NW_013535492.1 | 2832430 | G | 2 | G/C | rc\|pop | GCG | CGC | 0.05023 | intronic protein CIP2A homolog |
| NW_013535386.1 | 3083373 | G | 2 | G/A | pop | GGG | AAA | 0.05948 | intronic teneurin-3 (downstream of CYP4G cluster) |
| NW_013535423.1 | 1590364 | A | 2 | A/C | pop | AAA | CCC | 0.05948 | intronic neuropeptide F-like |
| NW_013535319.1 | 2357306 | G | 2 | G/A | pop | GGG | AAA | 0.06952 | intronic nuclear migration protein nudC |
| NW_013535323.1 | 7237902 | T | 2 | T/A | pop | TTT | AAA | 0.06952 | intergenic |
| NW_013535332.1 | 4322388 | A | 2 | A/T | pop | AAA | TTT | 0.06952 | intronic protein yellow-like |
| NW_013535362.1 | 4099150 | T | 2 | T/C | rc\|pop | CCT | TTC | 0.06952 | at intron-exon boundary uncharacterized LOC10612973 |
| NW_013535378.1 | 1393893 | T | 2 | T/C | pop | TTT | CCC | 0.06952 | intergenic |
| NW_013535378.1 | 874667 | T | 2 | T/C | pop | TTT | CCC | 0.06952 | intergenic |

**Table S5.** Gene expression data in transcripts per million mapped reads (TMM) from *A. transitella* midgut tissue of the susceptible (ALM) and resistant (R347) strains. Caterpillars used in this study were fed on standard diets (C) or diets containing bifenthrin (B). (Demkovich et al., 2019). The data is part of a full transcriptome dataset (NCBI SRA PRJNA548705)

| SCAFFOLD | START | END | STRAND | GENE_ID | RNA_id | ALM_B1 | ALM_B2 | ALM_B3 | ALM_C1 | ALM_C2 | ALM_C3 | R_B1 | R_B2 | R_B3 | R_C1 | R_C2 | R_C3 | functional annotation |
|---|---|---|---|---|---|---|---|---|---|---|---|---|---|---|---|---|---|---|
| NW_013535362.1 | 2922570 | 2976883 | + | gene3366 | rna4402 | 21.198 | 19.644 | 19.95 | 26.068 | 22.939 | 20.169 | 24.703 | 25.524 | 24.073 | 22.457 | 23.891 | 19.972 | dynein heavy chain, cytoplasmic (LOC106129619), mRNA |
| NW_013535362.1 | 2977372 | 2991119 | - | gene3367 | rna4403 | 13.341 | 11.162 | 12.851 | 13.071 | 11.069 | 11.717 | 14.156 | 12.192 | 15.585 | 15.498 | 11.538 | 11.719 | ADAM 17-like protease (LOC106129617), mRNA |
| NW_013535362.1 | 2996162 | 2999913 | - | gene3368 | rna4404 | 1157.374 | 1170.911 | 1380.37 | 845.41 | 947.265 | 963.473 | 773.743 | 928.322 | 926.916 | 941.836 | 800.146 | 925.398 | polyubiquitin-B (LOC106129618), mRNA |
| NW_013535362.1 | 3002489 | 3003978 | - | gene3369 | rna4405 | 61.76 | 62.52 | 65.665 | 67.124 | 56.987 | 60.824 | 59.531 | 42.463 | 44.688 | 52.877 | 59.556 | 46.742 | cytochrome c oxidase assembly protein COX16 homolog, mitochondrial ( |
| NW_013535362.1 | 3012239 | 3045757 | + | gene3370 | rna4407 | 13.136 | 12.626 | 13.481 | 13.886 | 11.999 | 10.649 | 11.114 | 11.578 | 14.678 | 10.127 | 9.376 | 11.568 | tyrosine-protein kinase CSK (LOC106129714), transcript variant X2, mRN |
| NW_013535362.1 | 3044389 | 3048122 | - | gene3371 | rna4408 | 0 | 0 | 0 | 0 | 0 | 0.37 | 0 | 0 | 0 | 0 | 0 | 0 | uncharacterized LOC106129572 (LOC106129572), mRNA |
| NW_013535362.1 | 3049409 | 3053897 | - | gene3372 | rna4409 | 5.537 | 4.442 | 5.773 | 4.339 | 4.65 | 4.19 | 4.506 | 5.388 | 5.05 | 4.346 | 5.297 | 4.265 | DET1 homolog (LOC106129605), mRNA |
| NW_013535362.1 | 3055117 | 3067241 | + | gene3373 | rna4410 | 3.627 | 3.144 | 4.009 | 3.556 | 4.74 | 4.036 | 2.949 | 2.752 | 3.337 | 2.489 | 2.447 | 2.781 | cyclin-dependent kinase 7 (LOC106129620), mRNA |
| NW_013535362.1 | 3067370 | 3078948 | - | gene3374 | rna4411 | 0.076 | 0.031 | 0.064 | 0 | 0 | 0.062 | 0.33 | 0.472 | 1.23 | 0.536 | 1.523 | 0.942 | tubulin polyglutamylase TTLL13-like (LOC106129573), mRNA |
| NW_013535362.1 | 3079983 | 3096923 | + | gene3375 | rna4412 | 60.637 | 59.129 | 64.392 | 68.098 | 61.366 | 54.961 | 52.067 | 50.746 | 54.98 | 51.307 | 57.925 | 60.033 | ubiquilin-1-like (LOC106129708), mRNA |
| NW_013535362.1 | 3097871 | 3099727 | + | gene3376 | rna4413 | 2.569 | 2.7 | 1.828 | 2.572 | 3.09 | 2.66 | 4.516 | 3.304 | 5.292 | 2.91 | 2.938 | 2.63 | GPI mannosyltransferase 2 (LOC106129667), transcript variant X2, mRNA |
| NW_013535362.1 | 3101631 | 3107700 | - | gene3377 | rna4415 | 18.759 | 19.129 | 29.187 | 18.384 | 21.319 | 21.093 | 19.074 | 25.043 | 23.569 | 24.917 | 21.395 | 22.984 | methyl-CpG-binding domain protein 3 (LOC106129668), transcript varian |
| NW_013535362.1 | 3128643 | 3157699 | + | gene3378 | rna4417 | 12.671 | 14.697 | 11.653 | 11.261 | 13.809 | 14.777 | 13.414 | 13.786 | 14.214 | 9.055 | 10.408 | 9.515 | phospholipid-transporting ATPase ID (LOC106129670), mRNA |
| NW_013535362.1 | 3159383 | 3164900 | + | gene3379 | rna4418 | 4.879 | 5.679 | 4.116 | 4.668 | 4.8 | 5.833 | 5.805 | 4.791 | 5.171 | 4.987 | 4.59 | 4.14 | uncharacterized LOC106129757 (LOC106129757), mRNA |
| NW_013535362.1 | 3164948 | 3193977 | - | gene3380 | rna4420 | 7.599 | 6.988 | 7.633 | 8.594 | 8.2 | 7.425 | 8.558 | 9.316 | 10.232 | 9.046 | 7.558 | 9.471 | unconventional myosin-IXa-like (LOC106129709), transcript variant X1, m |
| NW_013535362.1 | 3197609 | 3199024 | - | gene3381 | rna4421 | 0.086 | 0.165 | 0.086 | 0.286 | 0 | 0 | 0 | 0 | 0 | 0.392 | 0 | 0 | Krueppel-like factor 9 (LOC106129644), mRNA |
| NW_013535362.1 | 3204704 | 3279423 | - | gene3382 | rna4423 | 35.661 | 41.639 | 41.396 | 36.092 | 43.048 | 39.023 | 30.405 | 30.164 | 32.279 | 38.222 | 37.454 | 41.366 | protein groucho (LOC106129687), transcript variant X3, mRNA |
| NW_013535362.1 | 3304902 | 3307354 | + | gene3383 | rna4425 | 14.118 | 14.038 | 15.107 | 15.887 | 10.959 | 14.161 | 7.568 | 8.265 | 7.651 | 9.544 | 10.732 | 14.482 | phosphomevalonate kinase (LOC106129627), mRNA |
| NW_013535362.1 | 3344559 | 3370779 | + | gene3384 | rna4427 | 0.054 | 0.072 | 0.16 | 0.042 | 0.13 | 0.072 | 0.237 | 0.196 | 0.151 | 0.278 | 0.128 | 0.4 | sodium channel protein para (LOC106129645), transcript variant X2, mR |
| NW_013535362.1 | 3387986 | 3396394 | - | gene3385 | rna4428 | 635.126 | 942.27 | 790.564 | 741.602 | 963.904 | 1225.47 | 987.206 | 819.697 | 606.378 | 1120.149 | 1189.944 | 1363.85 | cytochrome P450 CYP6B54-55 |
| NW_013535362.1 | 3399243 | 3401314 | - | gene3386 | rna4429 | 1485.169 | 1056.611 | 1154.414 | 1002.488 | 692.72 | 1343.165 | 1244.519 | 1269.88 | 1720.282 | 931.067 | 635.679 | 984.551 | cytochrome P450 CYP6B56 |
| NW_013535362.1 | 3403909 | 3420607 | - | gene3387 | rna4430 | 32.272 | 29.92 | 32.554 | 35.404 | 32.358 | 33.015 | 36.478 | 32.925 | 35.948 | 40.979 | 39.842 | 43.135 | protein ariadne-1 (LOC106129696), transcript variant X1, mRNA |
| NW_013535362.1 | 3421473 | 3423241 | - | gene3388 | rna4432 | 34.409 | 35.661 | 40.295 | 30.778 | 31.048 | 36.548 | 29.446 | 30.565 | 33.74 | 33.809 | 29.503 | 32.846 | uncharacterized LOC106129721 (LOC106129721), mRNA |
| NW_013535362.1 | 3423369 | 3430390 | + | gene3389 | rna4433 | 39.007 | 40.381 | 36.692 | 47.057 | 44.187 | 52.414 | 42.324 | 35.294 | 41.553 | 33.915 | 38.456 | 27.667 | isoleucine--tRNA ligase, cytoplasmic (LOC106129722), mRNA |
| NW_013535362.1 | 3433176 | 3489393 | - | gene3390 | rna4434 | 16.255 | 14.924 | 16.593 | 17.178 | 17.539 | 14.202 | 16.527 | 20.43 | 20.736 | 22.658 | 19.41 | 23.437 | dmX-like protein 2 (LOC106129661), mRNA |
| NW_013535362.1 | 3476653 | 3481076 | + | gene3391 | rna4435 | 9.962 | 0 | 7.11 | 0 | 0 | 2.773 | 0 | 0 | 6.028 | 0 | 0 | 0 | uncharacterized LOC106129658 (LOC106129658), mRNA |
| NW_013535362.1 | 3479317 | 3481542 | - | gene3392 | rna4437 | 0 | 0 | 0 | 0 | 0.49 | 0 | 0 | 0 | 0 | 0 | 0 | 0 | uncharacterized LOC106129657 (LOC106129657), transcript variant X1, n |
| NW_013535362.1 | 3490744 | 3524106 | + | gene3393 | rna4438 | 20.756 | 20.469 | 28.192 | 19.475 | 20.209 | 19.296 | 21.487 | 24.981 | 30.474 | 21.538 | 22.525 | 34.055 | endophilin-A (LOC106129629), transcript variant X1, mRNA |
| NW_013535362.1 | 3525167 | 3528547 | - | gene3394 | rna4441 | 766.298 | 1001.831 | 1068.671 | 755.594 | 739.317 | 803.993 | 906.168 | 974.517 | 878.216 | 1018.051 | 918.148 | 1123.07 | polyubiquitin-C (LOC106129631), mRNA |
| NW_013535362.1 | 3529722 | 3548710 | + | gene3395 | rna4442 | 23.195 | 26.457 | 27.562 | 17.961 | 17.259 | 17.037 | 22.373 | 28.365 | 22.712 | 27.626 | 22.093 | 36.356 | ras GTPase-activating protein 1 (LOC106129632), mRNA |
| NW_013535362.1 | 3549071 | 3550177 | - | gene3396 | rna4443 | 52.758 | 53.481 | 48.313 | 56.857 | 55.657 | 56.501 | 41.87 | 43.461 | 24.647 | 50.034 | 55.438 | 42.593 | protein kish-A (LOC106129633), mRNA |
| NW_013535362.1 | 3550487 | 3554188 | + | gene3397 | rna4444 | 9.66 | 7.792 | 8.852 | 7.864 | 7.36 | 7.876 | 7.815 | 8.71 | 7.742 | 9.4 | 6.565 | 9.373 | protein prenyltransferase alpha subunit repeat-containing protein 1 (LOC |
| NW_013535362.1 | 3584697 | 3630468 | + | gene3398 | rna4445 | 0.151 | 0.278 | 0.15 | 0.191 | 0.31 | 0.175 | 0.32 | 0.338 | 0.343 | 0.211 | 0.128 | 0.24 | uncharacterized LOC106129575 (LOC106129575), mRNA |
| NW_013535362.1 | 3631722 | 3638019 | - | gene3399 | rna4446 | 1.23 | 0.258 | 0.631 | 0 | 0.23 | 0 | 0 | 0.338 | 2.046 | 0.364 | 0 | 0.586 | uncharacterized LOC106129590 (LOC106129590), mRNA |
| NW_013535362.1 | 3745239 | 3796518 | + | gene3400 | rna4447 | 0.173 | 0.227 | 0.192 | 0.191 | 0.34 | 0.246 | 0.33 | 0.249 | 0.393 | 0.163 | 0.216 | 0.373 | protein unc-13 homolog A (LOC106129589), mRNA |
| NW_013535362.1 | 3800374 | 3817393 | + | gene3401 | rna4448 | 0.572 | 1.68 | 1.967 | 2.413 | 1.69 | 1.89 | 2.175 | 1.648 | 0.938 | 1.254 | 2.29 | 2.754 | actin-binding Rho-activating protein-like (LOC106129626), mRNA |
| NW_013535362.1 | 3826528 | 3849030 | - | gene3402 | rna4449 | 0.259 | 0.32 | 0.075 | 0.064 | 0 | 0.082 | 0.093 | 0.071 | 0.171 | 0.077 | 0.167 | 0.444 | gonadotropin-releasing hormone II receptor-like (LOC106129576), mRNA |
| NW_013535362.1 | 3852565 | 4034421 | - | gene3403 | rna4450 | 0.27 | 0.216 | 0.46 | 0.402 | 0.35 | 0.349 | 0.536 | 0.579 | 0.706 | 0.316 | 0.383 | 0.453 | small conductance calcium-activated potassium channel protein (LOC106 |
| NW_013535362.1 | 4090436 | 4108229 | - | gene3404 | rna4451 | 7.966 | 5.741 | 10.05 | 8.541 | 7.38 | 8.113 | 7.434 | 8.38 | 11.724 | 7.878 | 8.865 | 7.463 | uncharacterized LOC106129735 (LOC106129735), mRNA |
| NW_013535362.1 | 4240437 | 4246388 | + | gene3405 | rna4452 | 0.065 | 0.134 | 0.064 | 0.106 | 0.18 | 0.062 | 0.144 | 0.169 | 0 | 0.373 | 0.138 | 0.178 | extensin-like (LOC106129597), mRNA |
| NW_013535362.1 | 4247890 | 4251407 | + | gene3406 | rna4453 | 0 | 0 | 0 | 0 | 0 | 0.031 | 0 | 0 | 0 | 0 | 0 | 0.027 | uncharacterized protein DDB_G0286591-like (LOC106129577), mRNA |

**Table S6.** Bifenthrin use as the active ingredient(s) under registered trade names from 2006 – 2017 in almond orchards. Usage intensity (*UI*) is equal to the pounds of bifenthrin applied for each product divided by the treated acres. Trade names which comprise "Other" include Bifenture® EC-CA, Capture® EC-Cal, Swagger®, Helena Bifenthrin® 2EC-Cal, Sniper® Helios, Bifen 2 Ag Gold-Cal, Brigade® 2EC, Fanfare® EC, SPECKoZ® Bifenthrin, Bifenture® LFC, and Bifen 25% EC.

| **Statewide** | **Brigade® WSB** | | | | **Fanfare™ 2EC** | | | | **Bifenture® 10DF** | | | | **Bifenture® EC** | | | |
|---|---|---|---|---|---|---|---|---|---|---|---|---|---|---|---|---|
| **Almond Use** | Applications | Pounds Bifenthrin | Treated Acres | *UI* | Applications | Pounds Bifenthrin | Treated Acres | *UI* | Applications | Pounds Bifenthrin | Treated Acres | *UI* | Applications | Pounds Bifenthrin | Treated Acres | *UI* |
| 2006 | 434 | 3,904 | 32,456 | 0.12 | —— | —— | —— | —— | —— | —— | —— | —— | —— | —— | —— | —— |
| 2007 | 1,398 | 9,979 | 96,946 | 0.10 | —— | —— | —— | —— | —— | —— | —— | —— | —— | —— | —— | —— |
| 2008 | 1,310 | 10,403 | 103,107 | 0.10 | —— | —— | —— | —— | —— | —— | —— | —— | —— | —— | —— | —— |
| 2009 | 1,433 | 13,819 | 123,986 | 0.11 | —— | —— | —— | —— | —— | —— | —— | —— | —— | —— | —— | —— |
| 2010 | 1,101 | 10,364 | 91,170 | 0.11 | 836 | 16,112 | 93,979 | 0.17 | 300 | 2,035 | 20,287 | 0.10 | —— | —— | —— | —— |
| 2011 | 683 | 5,574 | 51,574 | 0.11 | 743 | 12,904 | 74,310 | 0.17 | 139 | 737 | 9,530 | 0.08 | 1,326 | 29,504 | 112,093 | 0.26 |
| 2012 | 709 | 5,852 | 52,406 | 0.11 | 925 | 15,900 | 90,989 | 0.17 | 113 | 1,061 | 9,865 | 0.11 | 2,023 | 34,263 | 170,186 | 0.20 |
| 2013 | 772 | 6,144 | 52,306 | 0.12 | 516 | 6,207 | 33,981 | 0.18 | 127 | 1,104 | 10,995 | 0.10 | 3,167 | 41,515 | 242,981 | 0.17 |
| 2014 | 673 | 4,728 | 42,858 | 0.11 | 80 | 1,016 | 4,146 | 0.25 | 67 | 449 | 4,216 | 0.11 | 2,795 | 38,500 | 230,815 | 0.17 |
| 2015 | 799 | 5,794 | 51,726 | 0.11 | 99 | 518 | 3,322 | 0.16 | 90 | 499 | 4,729 | 0.11 | 3,437 | 50,974 | 281,786 | 0.18 |
| 2016 | 618 | 4,633 | 41,363 | 0.11 | 165 | 1,660 | 8,651 | 0.19 | 38 | 256 | 1,721 | 0.15 | 2,771 | 39,664 | 221,537 | 0.18 |
| 2017 | 622 | 4,364 | 48,724 | 0.09 | 183 | 1,763 | 10,428 | 0.17 | 8 | 60 | 547 | 0.11 | 2,950 | 34,773 | 202,369 | 0.172 |
| | **Sniper®** | | | | **Athena®** | | | | **Brigadier®** | | | | **Hero® EW** | | | |
| | Applications | Pounds Bifenthrin | Treated Acres | *UI* | Applications | Pounds Bifenthrin | Treated Acres | *UI* | Applications | Pounds Bifenthrin | Treated Acres | *UI* | Applications | Pounds Bifenthrin | Treated Acres | *UI* |
| 2011 | 140 | 5,044 | 12,840 | 0.39 | 158 | 901 | 9,603 | 0.09 | 1 | 7 | 65 | 0.10 | 3 | 13 | 243 | 0.053 |
| 2012 | 291 | 3,834 | 21,232 | 0.18 | 229 | 1,389 | 14,499 | 0.10 | 1 | 29 | 285 | 0.10 | —— | —— | —— | —— |
| 2013 | 604 | 9,226 | 49,684 | 0.19 | 188 | 1,490 | 12,872 | 0.12 | 9 | 56 | 702 | 0.08 | 1 | 3 | 40 | 0.075 |
| 2014 | 565 | 9,959 | 52,676 | 0.19 | 239 | 1,606 | 14,275 | 0.11 | 25 | 144 | 2,275 | 0.06 | 73 | 176 | 3,641 | 0.05 |
| 2015 | 1,026 | 17,569 | 92,445 | 0.19 | 311 | 2,074 | 18,394 | 0.11 | 109 | 424 | 5,554 | 0.08 | 96 | 397 | 6,898 | 0.06 |
| 2016 | 986 | 15,204 | 80,879 | 0.19 | 130 | 519 | 4,882 | 0.11 | 109 | 446 | 6,060 | 0.07 | 20 | 68 | 1,166 | 0.06 |
| 2017 | 1,293 | 25,618 | 132,966 | 0.193 | 121 | 586 | 5,300 | 0.11 | 28 | 32 | 611 | 0.05 | 92 | 264 | 4,441 | 0.06 |
| | **Bifen 2 Ag Gold** | | | | **Fanfare® ES** | | | | **Aceto Bifenthrin 2EC** | | | | **Other** | | | |
| | Applications | Pounds Bifenthrin | Treated Acres | *UI* | Applications | Pounds Bifenthrin | Treated Acres | *UI* | Applications | Pounds Bifenthrin | Treated Acres | *UI* | Applications | Pounds Bifenthrin | Treated Acres | *UI* |
| 2007 | —— | —— | —— | —— | —— | —— | —— | —— | —— | —— | —— | —— | 3 | 0.89 | 9.5 | 0.09 |
| 2008 | —— | —— | —— | —— | —— | —— | —— | —— | —— | —— | —— | —— | 2 | 0.55 | 5.5 | 0.10 |
| 2009 | —— | —— | —— | —— | —— | —— | —— | —— | —— | —— | —— | —— | —— | —— | —— | —— |
| 2010 | —— | —— | —— | —— | —— | —— | —— | —— | —— | —— | —— | —— | —— | —— | —— | —— |
| 2011 | —— | —— | —— | —— | —— | —— | —— | —— | —— | —— | —— | —— | —— | —— | —— | —— |
| 2012 | —— | —— | —— | —— | —— | —— | —— | —— | —— | —— | —— | —— | 3 | 42 | 305 | |
| 2013 | —— | —— | —— | —— | 325 | 4,978 | 25,036 | 0.20 | —— | —— | —— | —— | 38 | 1,132 | 5,805 | 0.20 |
| 2014 | 878 | 6,716 | 48,885 | 0.14 | 105 | 1,251 | 6,618 | 0.19 | 303 | 4,138 | 22,062 | 0.19 | 13 | 33 | 276 | 0.12 |
| 2015 | 919 | 6,430 | 49,713 | 0.13 | 246 | 3,260 | 17,368 | 0.19 | 432 | 5,139 | 32,214 | 0.16 | 47 | 628 | 4,820 | 0.13 |
| 2016 | 969 | 6,648 | 50,174 | 0.133 | 410 | 4,156 | 28,788 | 0.14 | 666 | 8,282 | 47,522 | 0.17 | 62 | 370 | 2,785 | 0.133 |
| 2017 | 963 | 7,251 | 49,020 | 0.148 | 616 | 7,110 | 40,214 | 0.18 | 772 | 11,092 | 64,190 | 0.17 | 302 | 3,227 | 18,939 | 0.17 |

**Figure S1.** Snapshot of IGV genome browser showing a portion of the region in scaffold NW_013535362.1. **A**. Reads of the three sequenced populations are identical to the reference genome; **B**. Portion of the region upstream of the total sweep, where only the resistant genotype (R347) maintains identity with the genome while the susceptible lines ALM and FIG have accumulated polymorphisms. The same pattern is see downstream of the total sweep (not shown). Read coverage tracks display gray for same nucleotide and color for nucleotide changes.

A.

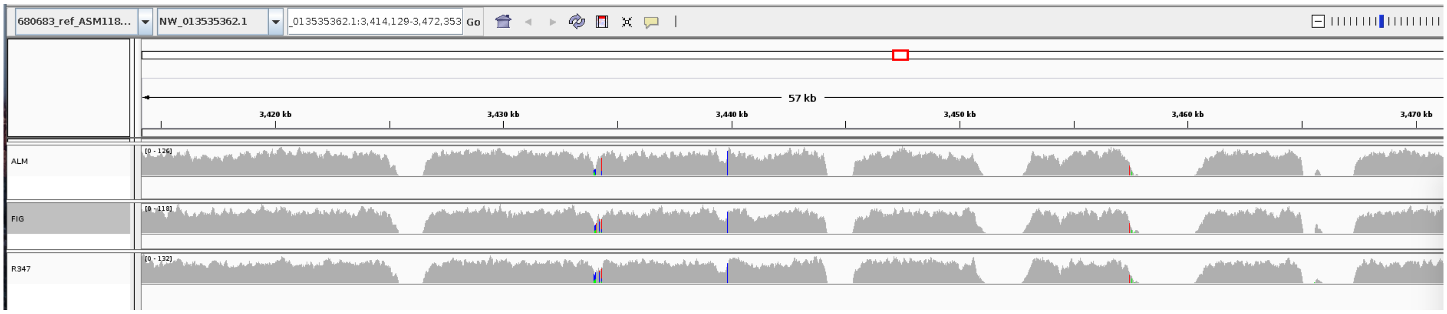


B.

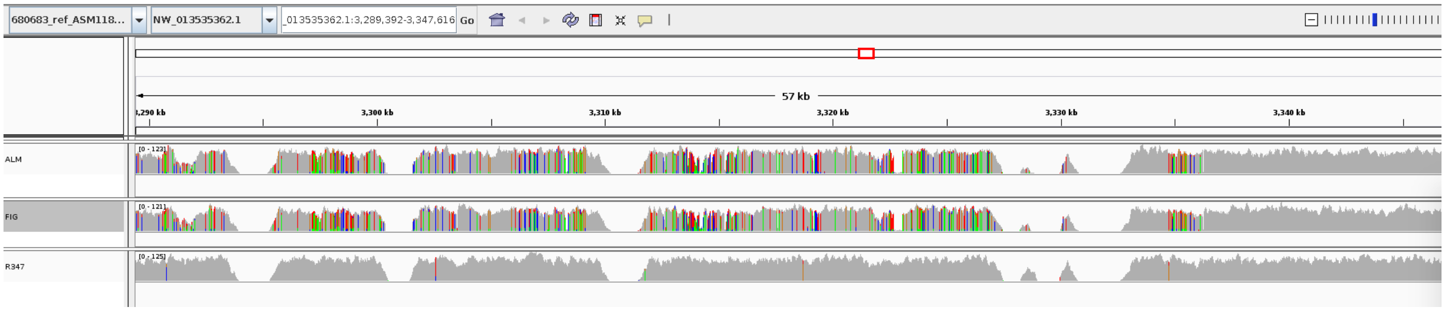

**Figure S2.** Sequence of the region flanking the position of the *kdr* mutation in the para gene in ten museum individuals of the SPIRL-1966 strain.

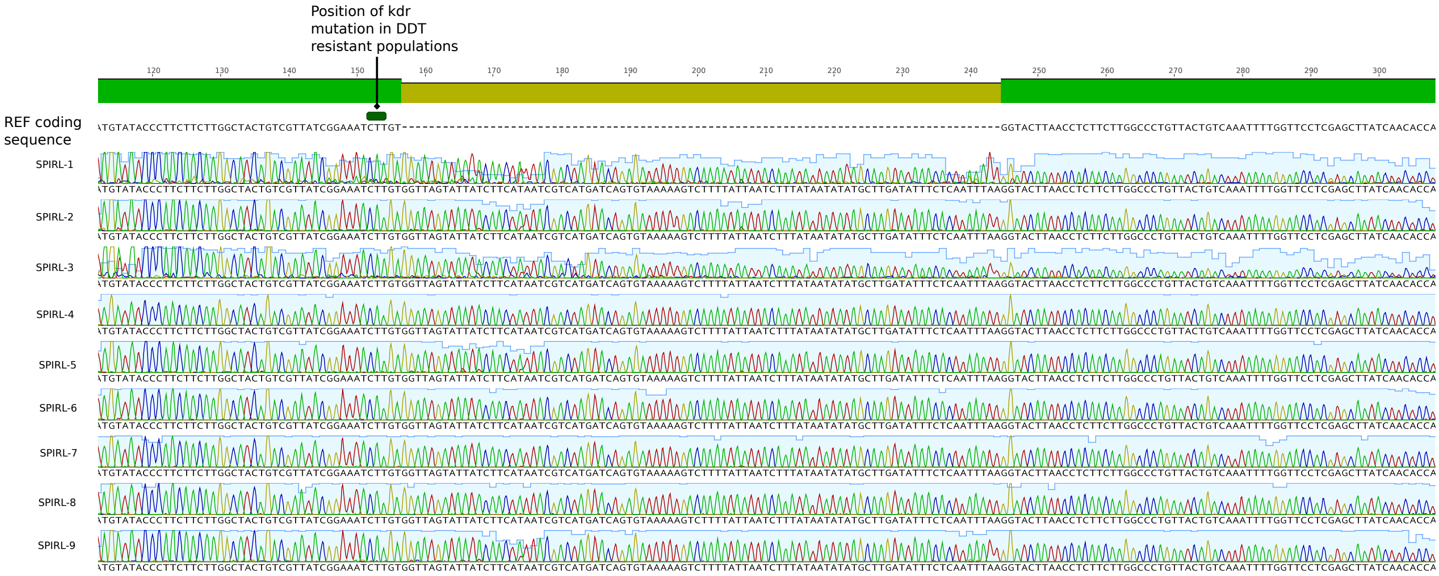

**Figure S3.** Nucleotide diversity specific for detoxificative gene families. **A**. Cytochrome P450s; **B.** Glutahtion-S-transferases; **C.** Carboxyl esterases **D.** ATP-binding cassette transporters.

A.

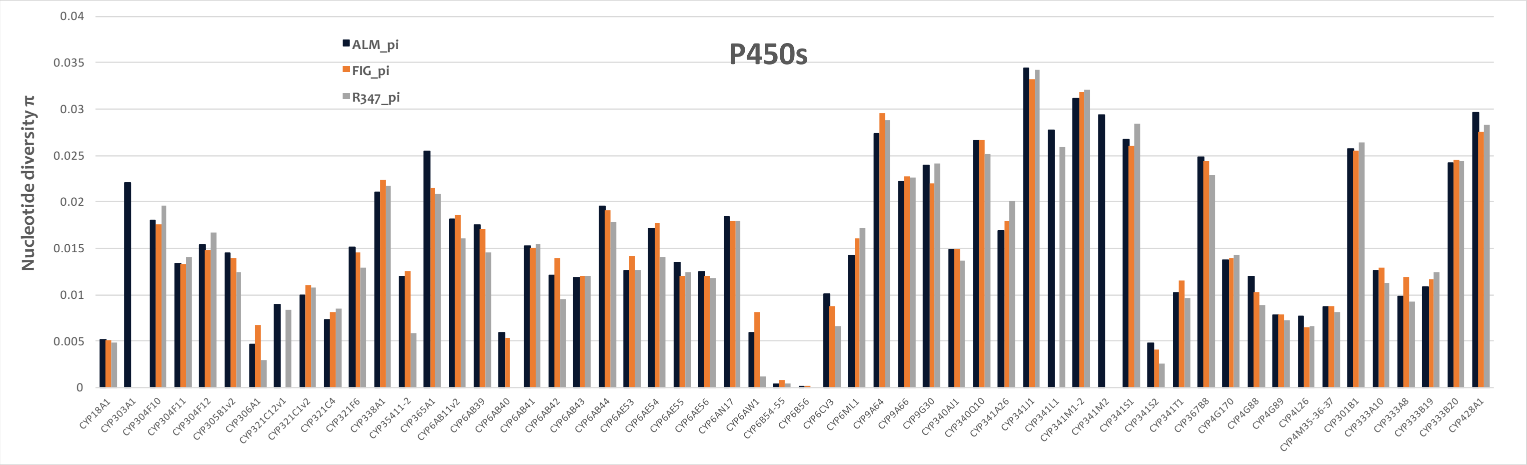


**B.**

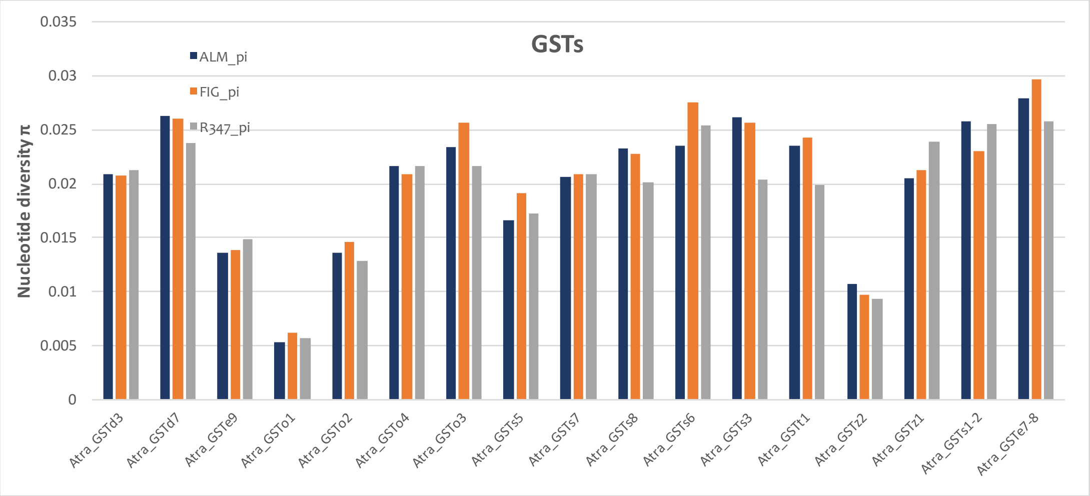


**C.**

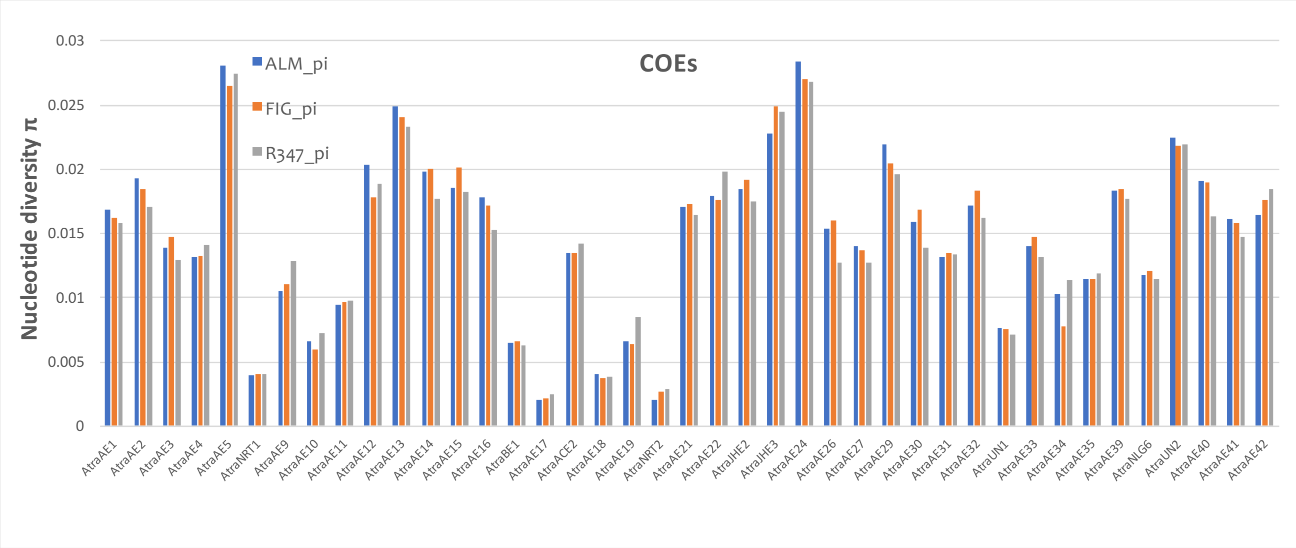
COEs
ALM_pi
FIG_pi
R347_pi
Nucleotide diversity π
0.03
0.025
0.02
0.015
0.01
0.005
0
AtraAE1
AtraAE2
AtraAE3
AtraAE4
AtraAE5
AtraNRT1
AtraAE9
AtraAE10
AtraAE11
AtraAE12
AtraAE13
AtraAE14
AtraAE15
AtraAE16
AtraBE1
AtraAE17
AtraACE2
AtraAE18
AtraAE19
AtraNRT2
AtraAE21
AtraAE22
AtraJHE2
AtraJHE3
AtraAE24
AtraAE26
AtraAE27
AtraAE29
AtraAE30
AtraAE31
AtraAE32
AtraUN1
AtraAE33
AtraAE34
AtraAE35
AtraAE39
AtraNLG6
AtraUN2
AtraAE40
AtraAE41
AtraAE42

**D.**

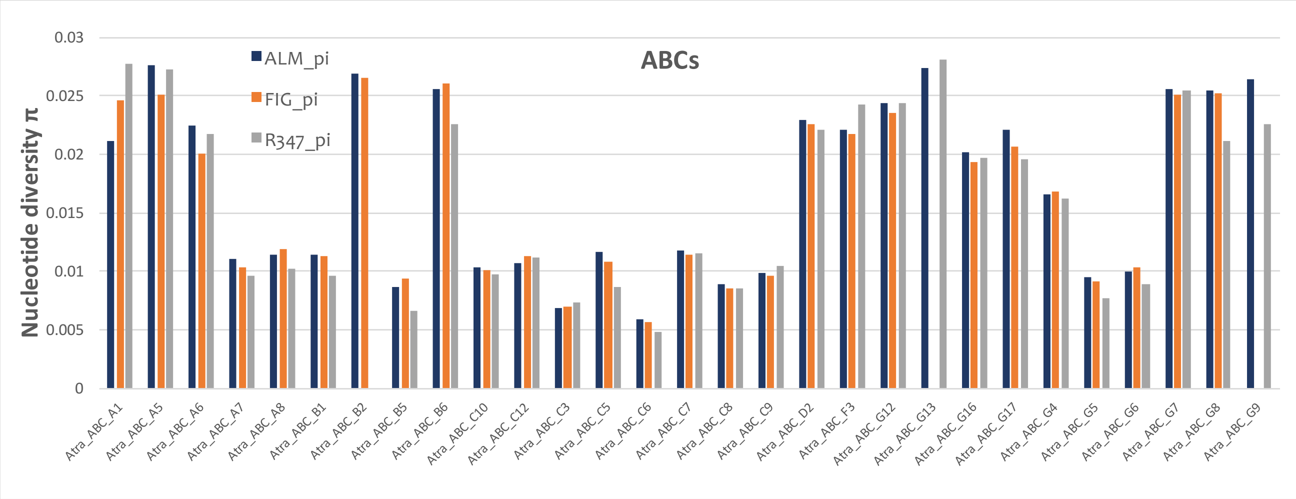
ABCs
ALM_pi
FIG_pi
R347_pi
Nucleotide diversity π
0.03
0.025
0.02
0.015
0.01
0.005
0
Atra_ABC_A1
Atra_ABC_A5
Atra_ABC_A6
Atra_ABC_A7
Atra_ABC_A8
Atra_ABC_B1
Atra_ABC_B2
Atra_ABC_B5
Atra_ABC_B6
Atra_ABC_C10
Atra_ABC_C12
Atra_ABC_C3
Atra_ABC_C5
Atra_ABC_C6
Atra_ABC_C7
Atra_ABC_C8
Atra_ABC_C9
Atra_ABC_D2
Atra_ABC_F3
Atra_ABC_G12
Atra_ABC_G13
Atra_ABC_G16
Atra_ABC_G17
Atra_ABC_G4
Atra_ABC_G5
Atra_ABC_G6
Atra_ABC_G7
Atra_ABC_G8
Atra_ABC_G9

**Figure S4.** Comparison of FST values obtained with Popoolation2 vs. Poolfstat. The screenshot shows the full reference genome coordinates (top). Tracks are pairwise FST values. Blue dots are values obtained with Popoolation2, and red dots are values obtained with Poolfstat**.**

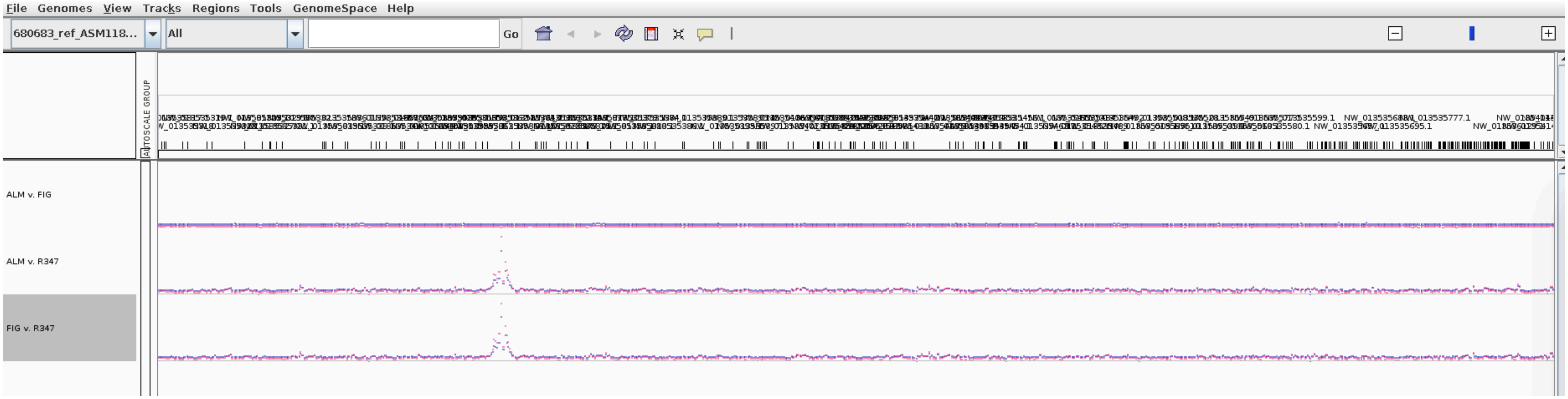

**Figure S5.** Bifenthrin use since its registration in 2006 versus all other pyrethroids reported in the DPR pesticide use records (cyfluthrin, beta-cyfluthrin, (S)-cypermethrin, deltamethrin, esfenvalerate, fenpropathrin, lambda-cyhalothrin, gamma-cyhalothrin, permethrin) in almond orchards from 2006 – 2017.

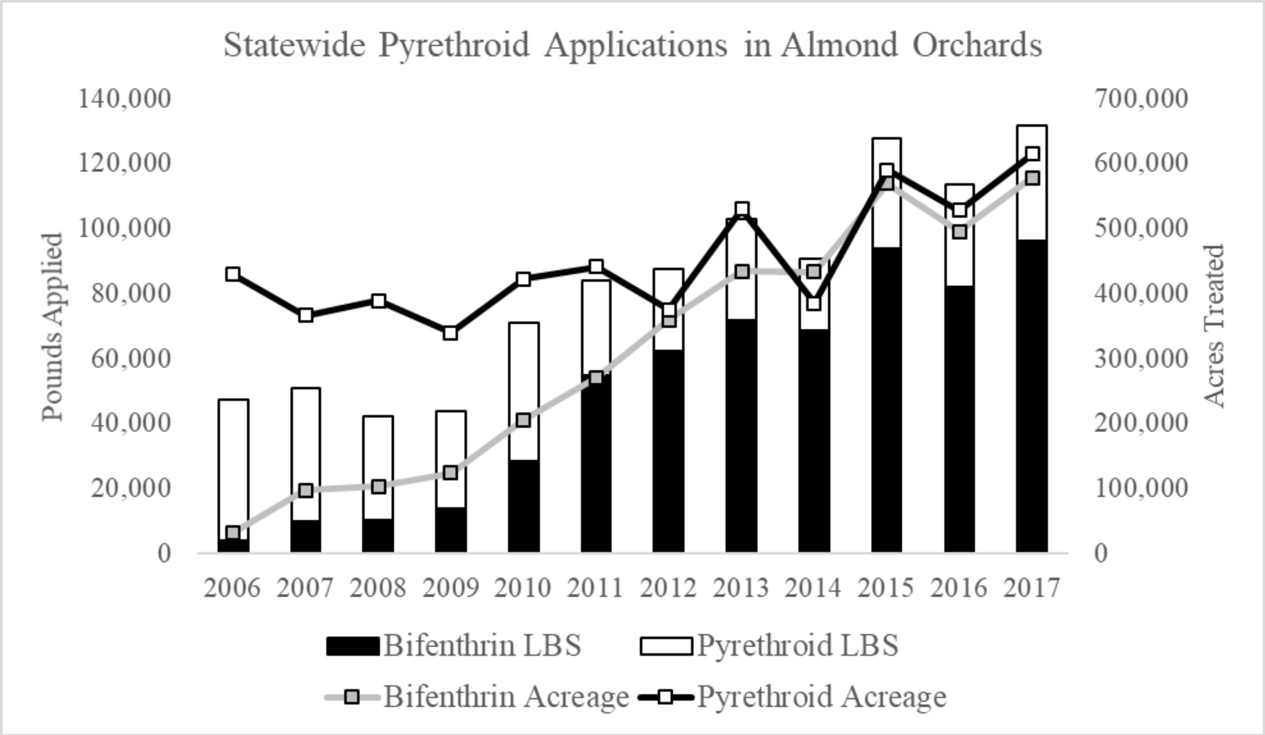
